# The insertion of a mitochondrial selfish element into the nuclear genome and its consequences

**DOI:** 10.1101/787044

**Authors:** Julien Y. Dutheil, Karin Münch, Klaas Schotanus, Eva H. Stukenbrock, Regine Kahmann

**Affiliations:** Max Planck Institute for Evolutionary Biology – Plön, Germany; Max Planck Institute for Terrestrial Microbiology – Marburg, Germany; Institute of Evolutionary Sciences CNRS – University of Montpellier – IRD – EPHE – Montpellier, France; Christian Albrechts University of Kiel – Kiel, Germany; Department of Molecular Genetics and Microbiology (MGM). Duke University Medical Center – Durham, NC, USA

**Keywords:** homing endonuclease, mitochondrion, intron, gene birth, gene transfer

## Abstract

Homing endonucleases (HE) are enzymes capable of cutting DNA at highly specific target sequences, the repair of the generated double-strand break resulting in the insertion of the HE-encoding gene (“homing” mechanism). HEs are present in all three domains of life and viruses; in eukaryotes, they are mostly found in the genomes of mitochondria and chloroplasts, as well as nuclear ribosomal RNAs. We here report the case of a HE that accidentally integrated into a telomeric region of the nuclear genome of the fungal maize pathogen *Ustilago maydis*. We show that the gene has a mitochondrial origin, but its original copy is absent from the *U. maydis* mitochondrial genome, suggesting a subsequent loss or a horizontal transfer from a different species. The telomeric HE underwent mutations in its active site and lost its original start codon. A potential other start codon was retained downstream, but we did not detect any significant transcription of the newly created open reading frame, suggesting that the inserted gene is not functional. Besides, the insertion site is located in a putative RecQ helicase gene, truncating the C-terminal domain of the protein. The truncated helicase is expressed during infection of the host, together with other homologous telomeric helicases. This unusual mutational event altered two genes: the integrated HE gene subsequently lost its homing activity, while its insertion created a truncated version of an existing gene, possibly altering its function. As the insertion is absent in other field isolates, suggesting that it is recent, the *U. maydis* 521 reference strain offers a snapshot of this singular mutational event.

## Introduction

The elucidation of the mechanisms at the origin of genetic variation is a longstanding goal of molecular evolutionary biology. Mutation accumulation experiments - together with comparative analysis of sequence data - are instrumental in studying the processes shaping genetic diversity at the molecular level (Kondrashov and Kondrashov 2010; Eyre-Walker and Keightley 2007). They revealed that the spectrum of mutations ranges from single nucleotide substitutions to large scale chromosomal rearrangements, and encompasses insertions, deletions, inversions, and duplication of genetic material of variable length (Lynch et al. 2008). Mutation events may result from intrinsic factors such as replication errors and repair of DNA damage. In some cases, however, mutations can be caused or favored by extrinsic factors, such as mutagenic environmental conditions or parasitic genome entities like viruses or selfish mobile elements. Such particular sequences, able to replicate and invade the host genome, may have various effects including inserting long stretches of DNA that do not encode any organismic function, but also disrupting, copying and moving parts of the genome sequence. These selfish element-mediated mutations can significantly contribute to the evolution of their host: first, the invasion of mobile elements creates “junk” DNA that can significantly increase the genome size (Lynch 2007), and some of this material can be ultimately domesticated and acquire a new function, beneficial to the host (Kaessmann 2010; Volff 2006). Second, the genome dynamics resulting from the activity of mobile elements can generate novelty by gene duplication (Ohta 2000; Dutheil et al. 2016) or serve as a mechanism of parasexuality and compensate for the reduced diversity in the absence of sexual reproduction (Dong et al. 2015; Möller and Stukenbrock 2017). Finally, control mechanisms (such as repeat-induced point mutations in fungi (Gladyshev 2017)) may also incidentally affect genetic diversity (Grandaubert et al. 2014).

Intron-borne homing endonuclease genes (HEG) constitute a class of selfish elements whose impact on genome evolution is less well documented. They encode a protein able to recognize a particular genomic DNA sequence and cut it (homing endonuclease, HE). The resulting double-strand break is subsequently repaired by recombination using the homologous sequence containing the HEG itself as a template, resulting in its insertion in the target location (Stoddard 2005). As the recognized sequence is typically large, its occurrence is rare and the insertion typically happens at a homologous position. In this process, a *heg*^+^ element containing the endonuclease gene converts a *heg*^-^ allele (devoid of HEG but harbouring the recognition sequence) to *heg*^+^, a mobility mechanism referred to as *homing* (Dujon et al. 1989). After the insertion, the host cell is homozygous *heg*^+^, and the HEG segregates at a higher frequency than the Mendelian rate (Goddard and Burt 1999). The open reading frame of the HEG is typically associated with a sequence capable of self-splicing, either at the RNA (group-I introns) or protein (inteins) level, avoiding disruption of functionality when inserted in a proteincoding gene (Chevalier and Stoddard 2001; Stoddard 2005). The dynamic of HEGs has been well described, and involves three stages: (i) conversion from *heg*^-^ to *heg*^+^ by homing activity, (ii) degeneration of the HEG leading to the loss of homing activity, but still protecting against a new insertion because the target is altered by the insertion event and (iii) loss of the HEG leading to the restoration of the *heg*^-^ allele (Gogarten and Hilario 2006; Barzel et al. 2011). This cycle leads to recurrent gains and losses of HEG at a given genomic position, and ultimately to the loss of the HEG at the population level unless new genes invade from other locations or by horizontal gene transfer (Gogarten and Hilario 2006).

HEGs are found in all domains of life as well as in the genomes of organelles, mitochondria and chloroplasts (Stoddard 2005; Lambowitz and Belfort 1993; Belfort and Roberts 1997). In several fungi, HEGs are residents of mitochondria. Here, we study the molecular evolution of a HEG from the fungus *Ustilago maydis*, which serves as a model for the elucidation of (i) fundamental biological processes like cell polarity, morphogenesis, organellar targeting, and (ii) the mechanisms allowing biotrophic fungi to colonize plants and cause disease (Steinberg and Perez-Martin 2008; Djamei and Kahmann 2012; Ast et al. 2013). *U. maydis* is the most well-studied representative of smut fungi, a large group of plant pathogens, because of the ease by which it can be manipulated both genetically and through reverse genetics approaches (Vollmeister et al. 2012). Besides, its compact, fully annotated genome comprises only 20.5 Mb and is mostly devoid of repetitive DNA (Kämper et al. 2006). The genome sequences of several related species, *Sporisorium reilianum, S. scitamineum*, and *U. hordei* causing head smut in corn, smut whip in sugarcane and covered smut in barley, respectively, provide a powerful resource for comparative studies (Schirawski et al. 2010; Laurie et al. 2012; Dutheil et al. 2016). Here, we report the case of a mitochondrial HEG that integrated into the nuclear genome of *U. maydis*. This singular mutation event created two new genes: first the original endonuclease activity of the integrated HEG was inactivated by a deletion in the active site, leading to a frameshift and a new open reading frame containing the DNA-binding domain of the HEG (Derbyshire et al. 1997). Second, the integration of the HEG occurred within another protein-coding gene, leading to its truncation.

## Methods

### 1. Analysis of codon usage and GC content

*Ustilago maydis* gene models (genome version 2.0) were retrieved from the MIPS database (Mewes et al. 2011). Mitochondrial genes were extracted from the *U*. maydis full mitochondrial genome (Genbank accession number: NC_008368.1). Within-group correspondence analysis of synonymous codon usage was performed using the ade4 package for R, following the procedure described in (Charif et al. 2005). The proportion of G and C nucleotides was computed along with the first 10 kb of *U. maydis* chromosome 9, using 300 bp windows slid by 1 bp. The corresponding R code is available as Supplementary File S1.

### 2. Strains, growth conditions and virulence assays

The *Escherichia coli* strains DH5α (Bethesda Research Laboratories) and TOP10 (Life Technologies, Carlsbad, CA, USA) were used for the cloning and amplification of plasmids. *U. maydis* strains 518 and 521 are the parents of FB1 and FB2 (Banuett and Herskowitz 1989). SG200 is a haploid solopathogenic strain derived from FB1 (Kämper et al. 2006). 10-1 is an uncharacterized haploid *U. maydis* strain isolated in the US and kindly provided by G. May. I2, O2, P2, S5, and T6 are haploid *U. maydis* strains collected in different parts of Mexico (Valverde et al. 2000). The haploid *S. reilianum* strains SRZ1 and SRZ2 as well as the solopathogenic strain JS161 derived from SRZ1 have been described (Schirawski et al. 2010). Deletion mutants were generated by gene replacement using a PCR-based approach and verified by Southern analysis (Kämper 2004).

pRS426Δum11064+11065 is a pRS426-derived plasmid containing the *UMAG_11064/ UMAG_11065* double deletion construct which consists of a hygromycin resistance cassette flanked by the left border of the *UMAG_11064* and right border of the *UMAG_11065* gene. The left border of *UMAG_11064* and the right border of *UMAG_11065* were PCR amplified from SG200 gDNA with primers um11064_lb_fw/um11064_lb_rv and um11065_rb_fw/um11065_rb_rv (Supplementary Table S7). The hygromycin resistance cassette was obtained from SfiI digested pHwtFRT (Khrunyk et al. 2010). The pRS426 EcoRI/XhoI backbone, both borders and the resistance cassette were assembled using yeast drag and drop cloning (Christianson et al. 1992). The fragment containing the deletion cassette was amplified from this plasmid using primers um11064_lb_fw and um11065_rb_rv (Supplementary Table S7), transformed into SG200 and transformants carrying a deletion of *UMAG_11064* and *UMAG_11065* were identified by southern analysis (Figure S3).

*U. maydis* strains were grown at 28°C in liquid YEPSL medium (0.4% yeast extract, 0.4% peptone, 2% sucrose) or on PD solid medium (2.4% Potato Dextrose broth, 2% agar). Stress assays were performed as described in (Krombach et al. 2018). Transformation and selection of *U. maydis* transformants followed published procedures (Kämper et al. 2006). To assess virulence, seven day old maize seedlings of the maize variety Early Golden Bantam (Urban Farmer, Westfield, Indiana, USA) were syringe-infected. At least three independent infections were carried out and disease symptoms were scored according to Kämper et al. (Kämper et al. 2006). Consistence of replicates was tested using a chi-squared test and p-values were computed using 1,000,000 permutations. As no significant difference between replicates was observed (p-value = 0.347 for the wildtype and p-value = 0.829 for the deletion strain), observation were pooled between all replicates for each strain before being compared.

### 3. Blast searches and gene alignment

We performed BlastN and BlastP (Altschul et al. 1990) searches using the (translated) sequence of *UMAG_11064* as a query using NCBI online blast tools. The non-redundant nucleotide and protein sequence databases were selected for BlastN and BlastP, respectively. Results were further processed with scripts using the NCBIXML module from BioPython (Cock et al. 2009). The Macse codon aligner (Ranwez et al. 2011) was used in order to infer the position of putative frameshifts in the upstream region of *UMAG_11064*. The alignment was depicted using the Boxshade software and was further manually annotated. The sequences of *U. maydis cox1* intron 6, as well as *S. reilianum cox1* introns 1 and 2 were used as query and searched against the protein non redundant database using NCBI BlastX, excluding environmental samples and model sequences. The *cox1* genes from *U. maydis* and *S. reilianum* were aligned and pairwise similarity was computed in non-overlapping 100 bp windows (Supplementary File S1). The gene structure, synteny and local pairwise similarity was depicted using the genoPlotR package for R (Guy et al. 2010).

### 4. Phylogeny estimation, estimation of dN/dS ratios and tests of positive selection

The nucleotide sequence of *UMAG_11064*, the first intron of the *cox1* gene of *S. reilianum*, and the eight non-redundant, most similar matches from BlastP (Table S2) were aligned using the Macse codon aligner (Ranwez et al. 2011) together with the unannotated but similar nucleotide sequences from *S. scitamineum, U. bromivora, T. indica* and *T. walkiri*, using the NCBI codon translation table 4 “mitochondrial mold”. Columns in the alignment were manually selected to discard ambiguously aligned regions and a phylogeny was inferred using PhyML (Guindon et al. 2010) with a General Time Reversible (GTR) model of nucleotide evolution and a 4-classes discrete gamma distribution of rates. The tree topology was inferred using the “best of nearest-neighbor-interchange (NNI) and subtree-pruning-regrafting (SPR)” option, and 100 non-parametric bootstrap replicates were obtained. The final tree was rooted using the midpoint method. Analyses were performed using the Seaview software (Gouy et al. 2010). For the positive selection analysis, the *S. scitamineum* and *U. bromivera* sequences were discarded as they contained multiple frameshifts. A phylogenetic tree was estimated using PhyML from the remaining species after translation using a Le and Gascuel model of protein evolution (Le and Gascuel 2008), and other parameters as for the nucleotide model. Nodes with bootstrap support values lower than 65% were collapsed. A branch model of codon evolution was fitted on the alignment and the inferred phylogenetic tree using PAML 4.9d (Yang 2007), keeping selected positions that may contain missing data. The F3X4 codon frequencies model was selected, and one dN/dS ratio was estimated per branch. A branch-site model (Zhang et al. 2005) was fitted by specifying the branch leading to the *UMAG_11064* gene as the “foreground” group, putatively evolving under positive selection. Test for selection was performed as suggested in the PAML manual, comparing to a model where the omega2 parameter is fixed to a value of 1. A similar test was conducted after excluding the two *Tilletia* sequences from the “background” branches, as they were found to have each a branch with dN/dS > 1.

### 5. Amplification of the *UMAG_11064* regions in several *U. maydis* strains

Amplification of DNA fragments via polymerase chain reaction (PCR) was done using the Phusion High Fidelity DNA_Polymerase (Thermo Fisher Scientific, Waltham, USA). The PCR reactions were set up in a 20 μl reaction volume using DNA templates indicated in the respective experiments and buffer recommended by the manufacturer containing a final concentration of 3% DMSO. The PCR programs used are represented by the following scheme: Initial denaturation – [denaturation – annealing – elongation] x number cycles – final elongation. *UMAG_11072* was amplified with primers um11072_ORF_fw x um11072_ORF_rv using 98 °C/3 m – [98 °C/10 s – 65 °C/30 s – 72 °C/45 s] x 30 cycles – 72 °C/10 m. *UMAG_11064* was amplified with primers um11064_ORF_fw x um11064_ORF_rv using 98 °C/3 m – [98 °C/10 s – 65 °C/30 s – 72 °C/45 s] x 30 cycles – 72 °C/10 m. The *cox1* exons 1+2 were amplified with primers cox1_ex1_rv x cox1_ex2_fw using 98 °C/3 m – [98 °C/10 s – 63 °C/30 s – 72 °C/90 s] x 33 cycles – 72 °C/10 m. cox1 exon 7 was amplified with primers cox1_ex7_fw X cox1_ex7_rv using 98 °C/3 m – [98 °C/10 s – 67 °C/30 s – 72 °C/60 s] x 30 cycles – 72 °C/10 m. Parts of the genomic region containing *UMAG_11064, UMAG_11065* and *UMAG_11066* were amplified with primer pairs um11064_fw1 x um11064_rv1, um11064_fw1 x um11064_rv2; and um11064_ fw2 x um11064_rv2 using 98 °C/3 m – [98 °C/10 s – 65 °C/30 s – 72 °C/150 s] x 32 cycles – 72 °C/10 m. The list of all primer sequences is provided in Supplementary Table S7. PCR results are shown in Figures S1 and S2.

### 6. History of the *UMAG_11065* family

The sequence of the *UMAG_11065* protein was used as a query for a search against several smut fungi (*U. maydis, U. hordei, S. reilianum, S. scitamineum, Melanopsichum pennsylvanicum, Pseudozyma flocculosa)* complete proteome using BlastP (Altschul et al. 1990). The search finds 17 hits within the *U. maydis* genome with an E-value below 0.0001, as well as two genes in *S. scitamineum* (*SPSC_04622* and *SPSC_05783*) and two genes in *P. flocculosa* (*PFL1_06135* and *PFL1_02192*). Using NCBI BlastP, we found several sequences from *Fusarium oxyparum* with high similarity. We selected the sequence *FOXG_04692* as a representative and added it to the data set. The Guidance web server with the GUIDANCE2 algorithm was then used to align the protein sequences and assess the quality of the resulting alignment. Default options from the server were kept, selecting the MAFFT aligner (Katoh et al. 2002). Several sequences appeared to be of shallow alignment quality and were discarded. The remaining sequences were realigned using the same protocol. Four iterations were performed until the final alignment had a quality good enough for phylogenetic inference. The final alignment contained 14 sequences and had a global score of 0.79. These 14 alignable sequences contained 13 *U. maydis* sequences (including *UMAG_11065*), and the *F. oxysporum* gene, other sequences from smut genomes were too divergent to be unambiguously aligned. Using Guidance, we further masked columns in the alignment with a score below 0.93 (a maximum of one position out of 14 in the column was allowed to be uncertain).

A phylogenetic analysis was conducted using the program Seaview 4 (Gouy et al. 2010). First, a site selection was performed in order to filter regions with too many gaps, leaving 506 sites. Second, a phylogenetic tree was built using PhyML within Seaview (Guindon et al. 2010) (Le and Gascuel protein substitution model (Le and Gascuel 2008) with a four-classes discretized gamma distribution of rates, the best tree of Nearest Neigbour Interchange (NNI) and Subtree Pruning and Regrafting (SPR) topological searches was kept). Support values were computed using the approximate likelihood ratio test (aLRT) method (Anisimova and Gascuel 2006).

A test for positive selection was conducted using a combination of branch and branch-site models using PAML (Yang 2007). The final GUIDANCE alignment was used, realigned using the Macse codon aligner (Ranwez et al. 2011), and ambiguously aligned sites and shorter sequences were manually filtered. The final alignment contained the following sequences: *UMAG_03394, UMAG_11065, UMAG_04486, UMAG_06506, UMAG_04094, UMAG_10585, UMAG_06474, UMAG_10980, UMAG_05977, FOXG_04692*. We used the PhyML software with the same options as described above to reconstruct a phylogenetic tree with this subset of sequences. The branch toward the *UMAG_11065* gene was used as a foreground group in the branch site model.

### 7. Gene expression

RNASeq normalized expression counts for the *UMAG_11064* and *UMAG_11065*, as well as of neighbouring genes and paralogs elsewhere in the genome, were extracted from the Gene Expression Omnibus data set GSE103876 (Lanver et al. 2018). Gene clustering based on expression profiles was conducted using a hierarchical clustering with an average linkage on a Canberra distance, suitable for expression counts, as implemented in the ‘dist’ and ‘hclust’ functions in R (R Core Team 2018). The resulting clustering tree was converted to a distance matrix and compared to the inferred phylogeny of the genes using a Mantel permutation test, as implemented in the ‘ape’ package for R (Paradis et al. 2004). Differences in expression between time points were assessed by fitting the linear model “expression ~ time * gene”, testing the effect of time while controlling for interaction with the “gene” variable. Residuals were normalized using a Box-Cox transform as implemented in the MASS package for R. Tukey’s posthoc comparisons were conducted on the resulting model, allowing for a 5% false discovery rate.

## Results

We report the analysis of the nuclear gene *UMAG_11064* from the smut fungus *U. maydis*, which was identified as an outlier in a whole-genome analysis of codon usage. We first provide evidence that the gene is a former HEG and then reconstruct the molecular events that led to its insertion in the nuclear genome using comparative sequence analysis. Finally, we assess the phenotypic impact of the insertion event.

### 1 The *UMAG_11064* nuclear gene has a mitochondrial codon usage

We studied the synonymous codon usage in protein-coding genes of the smut fungus *U. maydis*, using within-group correspondence analysis. As opposed to other methods, within-group correspondence analysis allows the comparison of codon usage while adequately taking into account confounding factors such as variation in amino-acid usage (Perrière and Thioulouse 2002). We report a distinct synonymous codon usage for nuclear genes and mitochondrial genes (Figure 1A), with the notable exception of the nuclear gene *UMAG_11064*, which displays a typical mitochondrial codon usage. The *UMAG_11064* gene is located in the telomeric region of chromosome 9, with no further downstream annotated gene (Figure 1B). It displays a low GC content of 30%, which contrasts with the GC content of the flanking regions (50%) and the rather homogeneous composition of the genome sequence of *U. maydis* as a whole. It is, however, in the compositional range of the mitochondrial genome (Figure 1B). Altogether, the synonymous codon usage and GC content of *UMAG_11064* suggest a mitochondrial origin.

**Figure 1.**
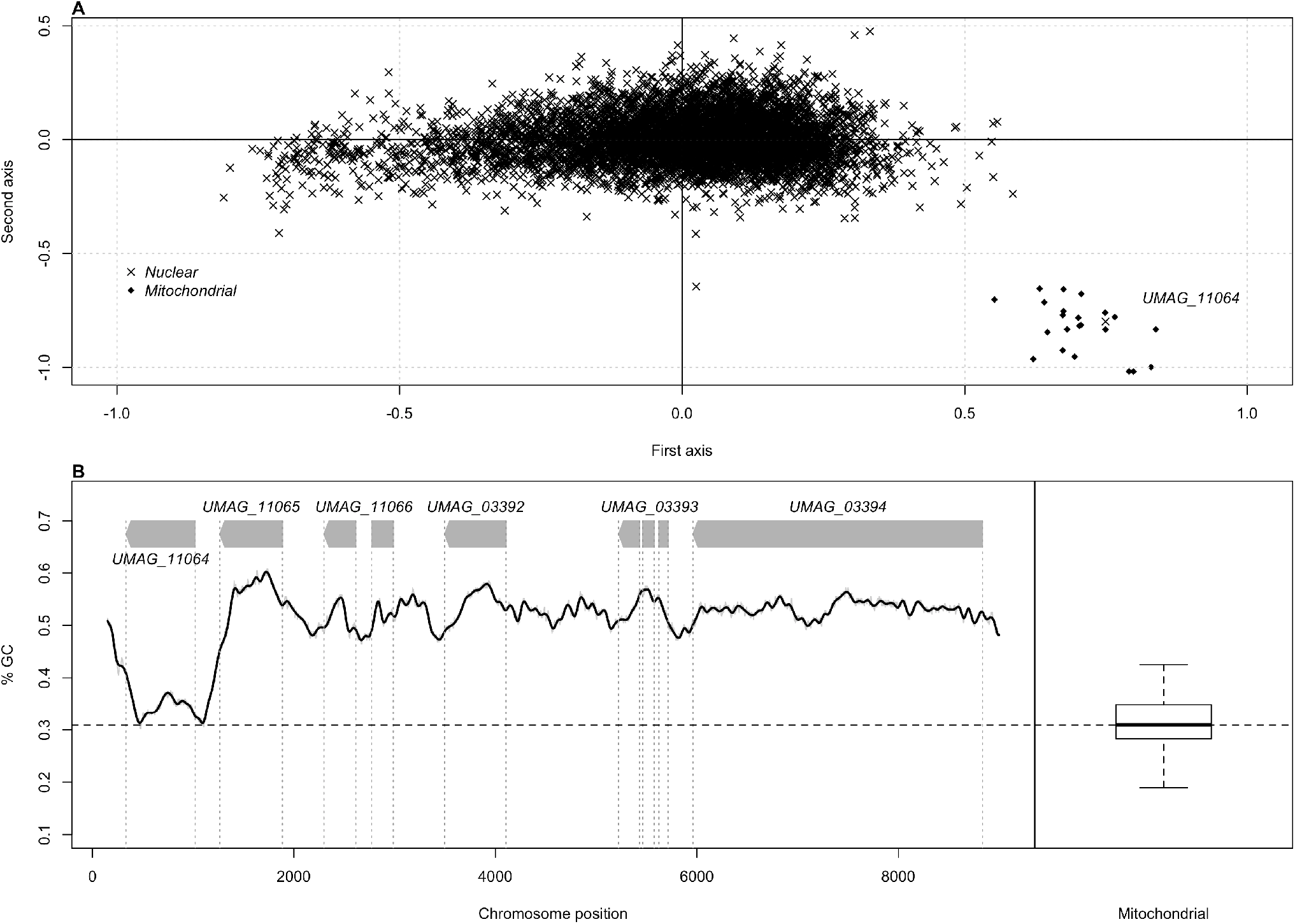
*Identification of the* UMAG_11064 *gene. A) Within-group correspondence analysis of* U. maydis *codon usage. Each gene is represented according to its coordinates along the first two principal factors. The genomic origin of each gene is indicated by a cross for nuclear genes and a dot for mitochondrial genes. B) Genomic context of the gene* UMAG_11064. *GC content in 300 bp windows sliding by 1 bp, and distribution of GC content in 300 bp windows of mitochondrial genome of U. maydis. The dash line represents the median of the distribution*.

In order to confirm the chromosomal location of *UMAG_11064*, we amplified and sequenced three regions encompassing the gene using primers within the *UMAG_11064* gene and primers in adjacent chromosomal genes upstream and downstream of *UMAG_11064* (Figure S1). The sequences of the amplified segments were in full agreement with the genome sequence of *U. maydis* (Kämper et al. 2006), thereby ruling out possible assembly artefacts in this region. As both the GC content and synonymous codon usage of *UMAG_11064* are indistinguishable from the ones of mitochondrial genes and have not moved toward the nuclear equilibrium, the transfer of the gene to its nuclear position is likely to have occurred relatively recently.

### 2 The *UMAG_11064* gene contains parts of a former GIY-YIG homing endonuclease

To gain insight into the nature of the *UMAG_11064* gene, its predicted nucleotide sequence was searched against the NCBI non-redundant nucleotide sequence database. Surprisingly, the sequence of *UMAG_11064* has no match in the mitochondrial genome of *U. maydis* itself (GenBank entry NC_008368.1), but high similarity matches were found in the mitochondrial genome of three other smut fungi (Supplementary Table S1): *S. reilianum* (87% nucleotide identity)*, S. scitamineum* (79%), and *U. bromivora* (76%). Two other very similar sequences were found in the mitochondrial genome of two other smut fungi, *Tilletia indica* and *Tilletia walkeri* (69% nucleotide identity), as well as in mitochondrial genomes from other basidiomycetes (e.g. *Laccaria bicolor*, 72%) and ascomycetes (e.g. *Leptosphaeria maculans*, 69%, see Supplementary Table S1). The protein sequence of *UMAG_11064* shows high similarity with fungal HEGs, in particular of the so-called GIY-YIG family (Supplementary Table S2) (Stoddard 2005). The closest fully annotated protein sequence matching *UMAG_11064* corresponds to the GIY-YIG HEG located in intron 1 of the *cox1* gene of *Agaricus bisporus* (I-AbiIII-P, 54% nucleotide identity). The amino-acid sequence of UMAG_11064 matches the N-terminal part of this protein containing the DNA-binding domain of the HE (Derbyshire et al. 1997).

We performed a codon alignment of the *UMAG_11064* gene together with the most similar sequences identified by Blast, using the Macse codon aligner to infer sequence alignment in the presence of frameshifts (Ranwez et al. 2011). The sequences from *S. scitamineum* and *U. bromivora* appeared to have several frameshifts introducing stop codons, suggesting that these sequences are pseudogenes. We reconstructed the phylogeny of the nucleotide sequences after removing ambiguously aligned regions (Figure 2). The resulting tree shows that the closest relative of the *UMAG_11064* gene is the intronic sequence from *S. reilianum*.

**Figure 2.**
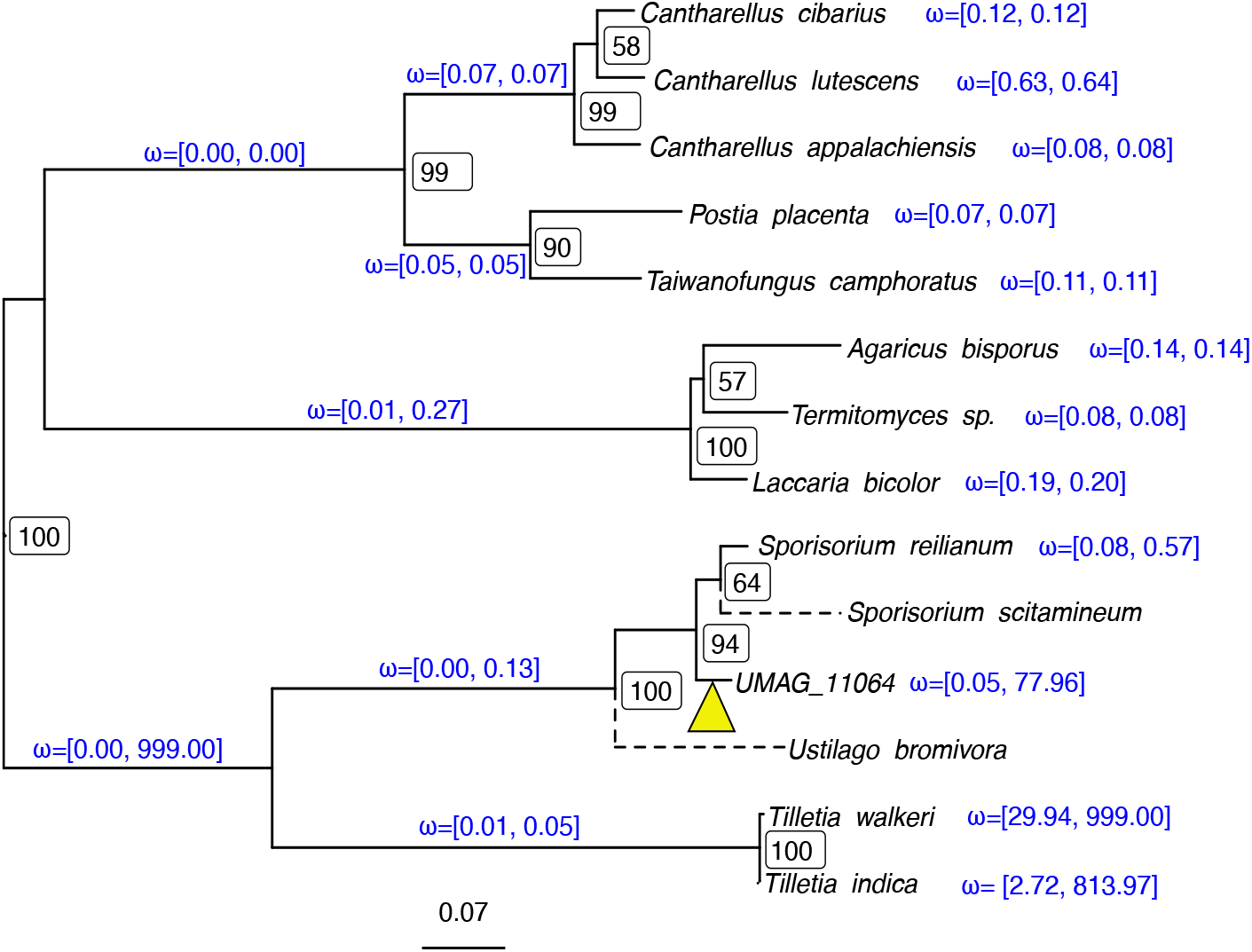
*Phylogeny of* UMAG_11064 *and its homologous sequences. Maximum likelihood tree inferred from nucleotide sequences. Node labels show bootstrap support values. Discontinuous branches indicate that the corresponding sequence is pseudogenized, with multiple frameshifts and insertions/deletions. Branch annotations show the minimum and maximum dN/dS ratio (ω) estimated from 10 independent runs of the codeml program. The yellow triangle indicates the supposed branch where the frameshift within the active domain of the ancestral HE occurred (See Figure 3)*.

As the GC profile of *UMAG_11064* suggests that the upstream region also has a mitochondrial origin (Figure 1B), we performed a codon alignment of the 5’ region with the full intron sequences of S*. reilianum, T. indica* and *T. walkeri* as well as the sequence of I-AbIII-P from *A. bisporus* in order to search for putative traces of the activity domain of the HE (Figure 3). We found that the intergenic region between *UMAG_11065* and *UMAG_11064* is similar to the activity domain of other GIY-YIG HE, and contains remnants of the former active site of the type GVY-YIG (Figure 3). Compared to I-AbiIII-P and homologous sequences in *Tilletia*, however, a frameshift mutation has occurred in the active site (a 7 bp deletion). The predicted gene model for *UMAG_11064* starts at a conserved methionine position, 14 amino-acids downstream of the former active site (Figure 3) and contains the *helix-turn-helix* DNA-binding domain of the original HE.

**Figure 3.**
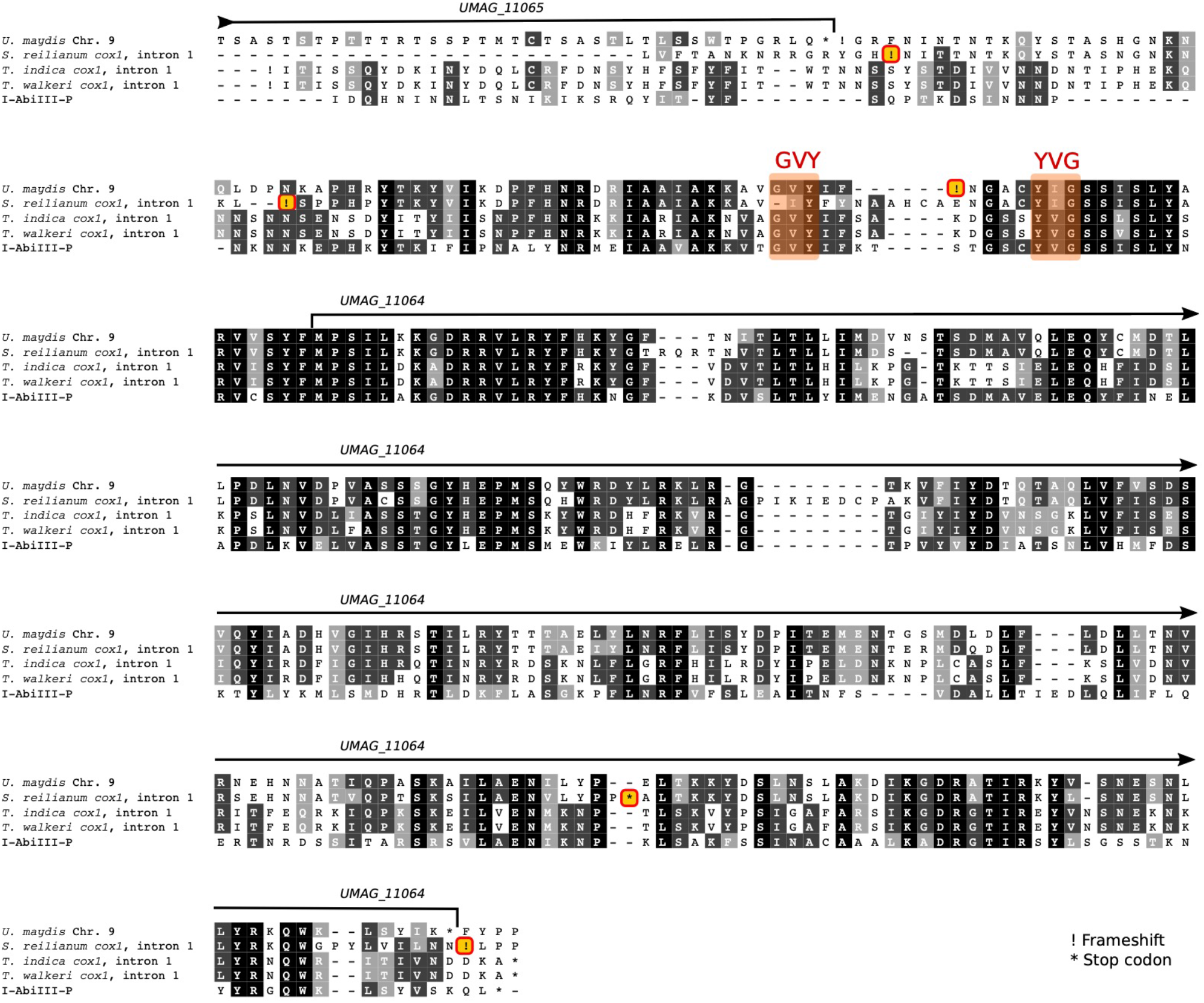
*Alignment of* UMAG_11064 *and its upstream sequence with intron 1 from the cox1 gene of* S. reilianum, T. indica *and* T. walkeri, *as well as the coding sequence of the* A. bisporus *HE. Shading indicates the level of amino-acid conservation, showing conserved residues (in black) and residues with similar biochemical properties (grayscale). Amino-acids noted as ‘X’ have incomplete codons due to frameshifts. Highlighted exclamation marks denote inferred frameshifts and * characters stop codons. The location of the active site of the HE (GVY-YVG) is highlighted*.

A branch model of codon sequence evolution was fitted to the codon sequence alignment of *UMAG_11064* and its identified homologs, with the exception of the putative pseudogenes from S. scitamineum and U. bromivora. The proteins appear to be evolving under purifying selection (dN/dS < 1) on most branches of the tree, with the exception of the branches leading to the two Tilletia sequences, as well as the branch leading to UMAG_11064 (Figure 2). We note, however, that the codeml program suffers from convergence issues on these particular branches, as witnessed by the variance in final estimates after 10 independent runs (Figure 2). Such convergence failures likely result from the branch model being an over-parametrized model, here fitted on relatively short sequences. To further assess whether the high dN/dS ratio measured in the *UMAG_11064* gene could be explained both by relaxed purifying selection or positive selection, we fitted a branch-site model allowing specifically for sites in the *UMAG_11064* gene to evolve under positive selection (foreground branch), which we contrasted with a null model where all sites evolve under purifying selection or neutral evolution. The likelihood ratio test was not significant (p-value equal to 0.1585), even after removing the *Tilletia* sequences (p-values equal to 0.2183), and does not reject the hypothesis that the *UMAG_11064* gene is evolving under a nearly neutral scenario. The higher dN/dS in the branch leading to *UMAG_11064* might, therefore, be the result of relaxed purifying selection.

Altogether, these results suggest that *UMAG_11064* is a former HE that inserted into the nuclear genome from the mitochondrion, was then inactivated by a deletion in its active site and acquired a new start codon, allowing it to code for a protein sequence with the former nucleotide binding domain of the HE.

### 3 The *UMAG_11064* gene is similar to an intronic mitochondrial sequence of *S. reilianum*

The closest homologous sequence of *UMAG_11064* was found in the first intron of the *cox1* gene of the smut fungus *S. reilianum* while this sequence was absent in the mitochondrial genome of *U. maydis*. The *cox1* genes of *S. reilianum* and *U. maydis* both have eight introns, of which only seven are homologous in position and sequence (Figure 4). *S. reilianum* has one extra intron in position 1, while *U. maydis* has one extra intron in position 6. In *U. maydis* all introns but the sixth one are reported to contain a HEG. A blast search of this intron’s sequence, however, revealed similarity with a homing endonuclease of type LAGLIDADG (Supplementary Table S4). In *S. reilianum*, intron 1 (the putative precursor of *UMAG_11064)* and intron 2 are not annotated as containing a HEG. Blast searches of the corresponding sequences, however, provided evidence for homology with a GIY-YIG HE (Supplementary Table S5) and a LAGLIDADG HE, respectively (Supplementary Table S6). Like many other species of fungi (Stone et al. 2018; Jalalzadeh et al. 2015; Pogoda et al. 2019), plants (Cho et al. 1998) and even animals (Fukami et al. 2007; Schuster et al. 2017), the *cox1* gene seems to be a hotspot of HEG-encoding introns in smut fungi.

**Figure 4.**
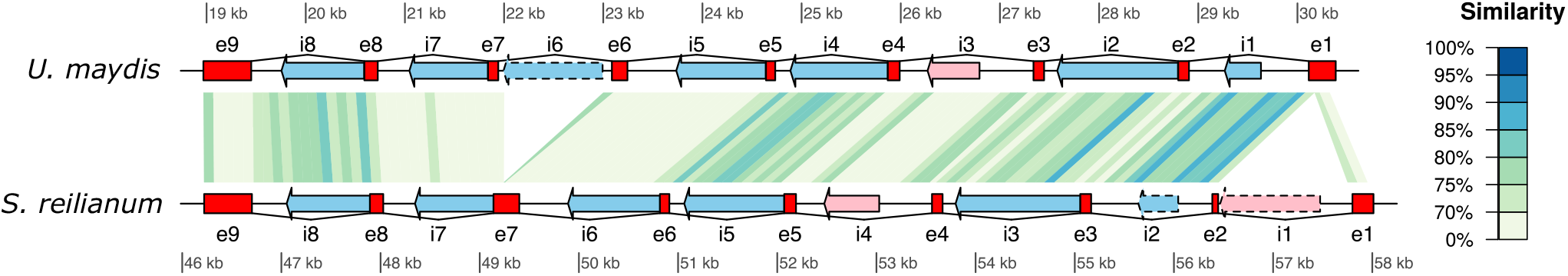
*Intron structure of the cox1 gene in* U. maydis *and* S. reilianum*. Annotated HEs are indicated. Red boxes depict cox1 exons, numbered from e1 to e9. Introns are represented by connecting lines and numbered i1 to i8. Arrows within introns show LAGLIDADG (light blue) and GIY-YIG HEs (pink). Dashed arrows correspond to HEGs inferred by blast search, while solid arrows correspond to the annotation from the GenBank files. Piecewise sequence similarity between* U. maydis *and* S. reilianum *is displayed with a color gradient*.

Lastly, no homolog of intron 1 in *S. reilianum* was detected in the mitochondrial genome of *U. maydis*. A closer inspection showed that the ORF could be aligned with related HEs in other fungi (Figure 3). This alignment revealed an insertion of four amino-acids, a deletion of the first glycine residue in the active site plus several frameshifts at the beginning of the gene, which suggests that this gene has been altered and might not encode a functional HE any longer.

### 4 *UMAG_11064* inserted into a gene encoding a RecQ helicase

In order to study the effect of the HEG insertion in the nuclear genome, we looked at the genomic environment of the *UMAG_11064* gene. Downstream of *UMAG_11064* are telomeric repeats, while the next upstream gene, *UMAG_11065*, is uncharacterized. A similarity search for *UMAG_11065* detected 13 paralogous sequences in the *U. maydis* genome (including one, *UMAG_12076*, on an unmapped contig), but only low-similarity matches in other sequenced smut fungi (see Methods). The closest non-smut related sequence comes from a gene from *Fusarium oxysporum*. We inferred the evolutionary relationships between the 14 genes by reconstructing a maximum likelihood phylogenetic tree, and found that the *UMAG_11065* gene is closely related to *UMAG_04486*, located on chromosome 14 (Figure 5 and Table 1). The *UMAG_04486* gene, however, is predicted to be almost six times as long as *UMAG_11065*. We note that the downstream region of *UMAG_11064* does not show any similarity with the 3’ part of the *UMAG_04486* gene, suggesting that the insertion of the HEG did not lead to the formation of an intron in the *UMAG_11065* gene, but rather to its truncation. A search for similar sequences of *UMAG_11065* and its relatives in public databases revealed homology with so-called RecQ helicases (Supplementary Table S3), enzymes known to be involved in DNA repair and telomere expansion (Singh et al. 2012). While this function is only predicted by homology, we note that all 12 chromosomal *recQ* related genes are located very close to telomeres in *U. maydis* (Table 1), suggesting a role of these genes in telomere maintenance (Sánchez-Alonso and Guzmán 1998). Lastly, we tested whether the truncation of *UMAG_11065* was followed by positively selected mutations in the remaining part of the gene. We inferred a dN/dS ratio equal to 0.342, which suggests that the *UMAG_11065* gene evolved mostly under purifying selection since divergence from the *UMAG_04486* gene. The insertion of UMAG_11064, therefore, was not followed by positive selection, or was too recent for sufficient positively selected substitutions to occur.

**Table 1.**
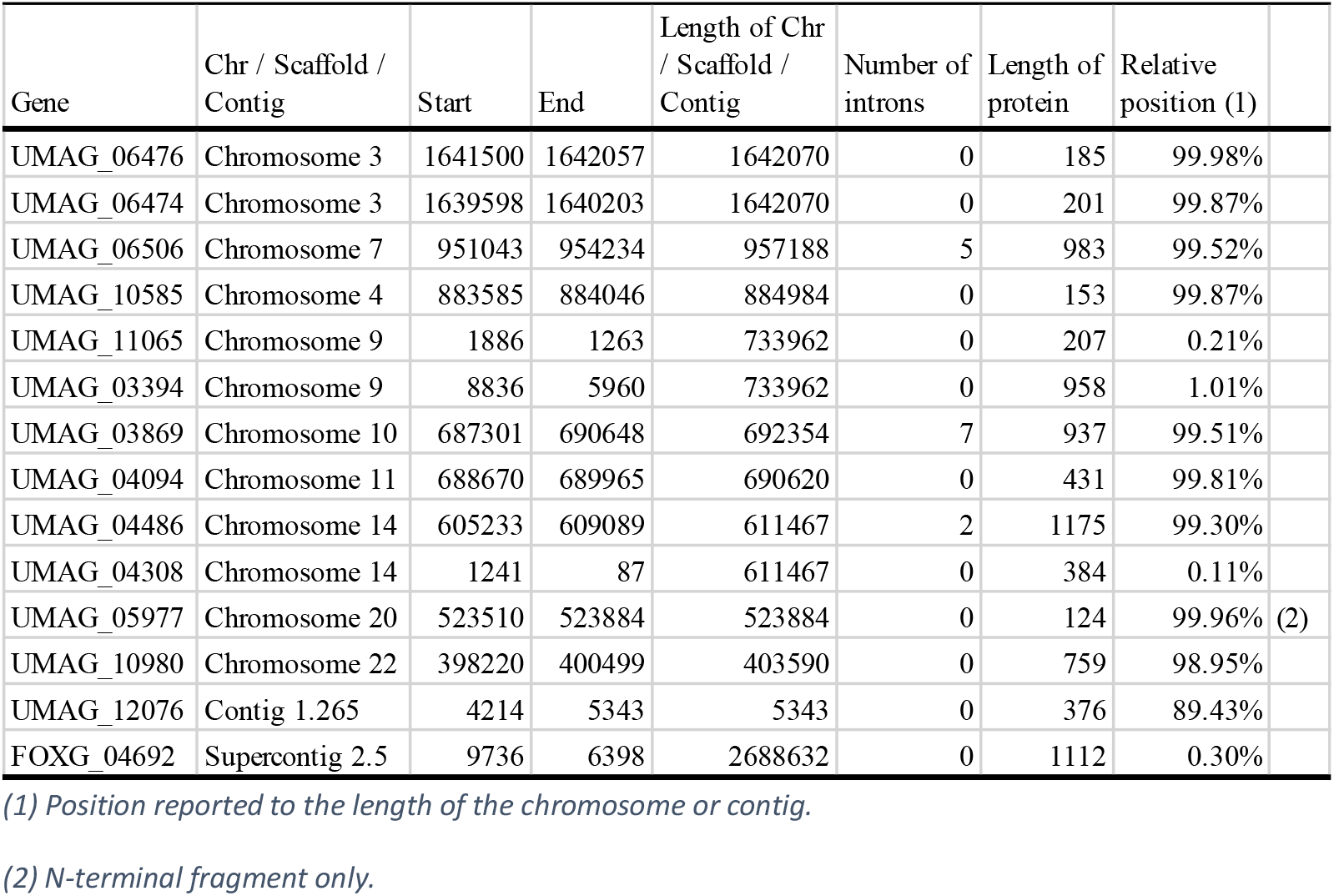
UMAG_11065 paralogs in U. maydis, together with a homolog from F. oxysporum for comparison.

**Figure 5.**
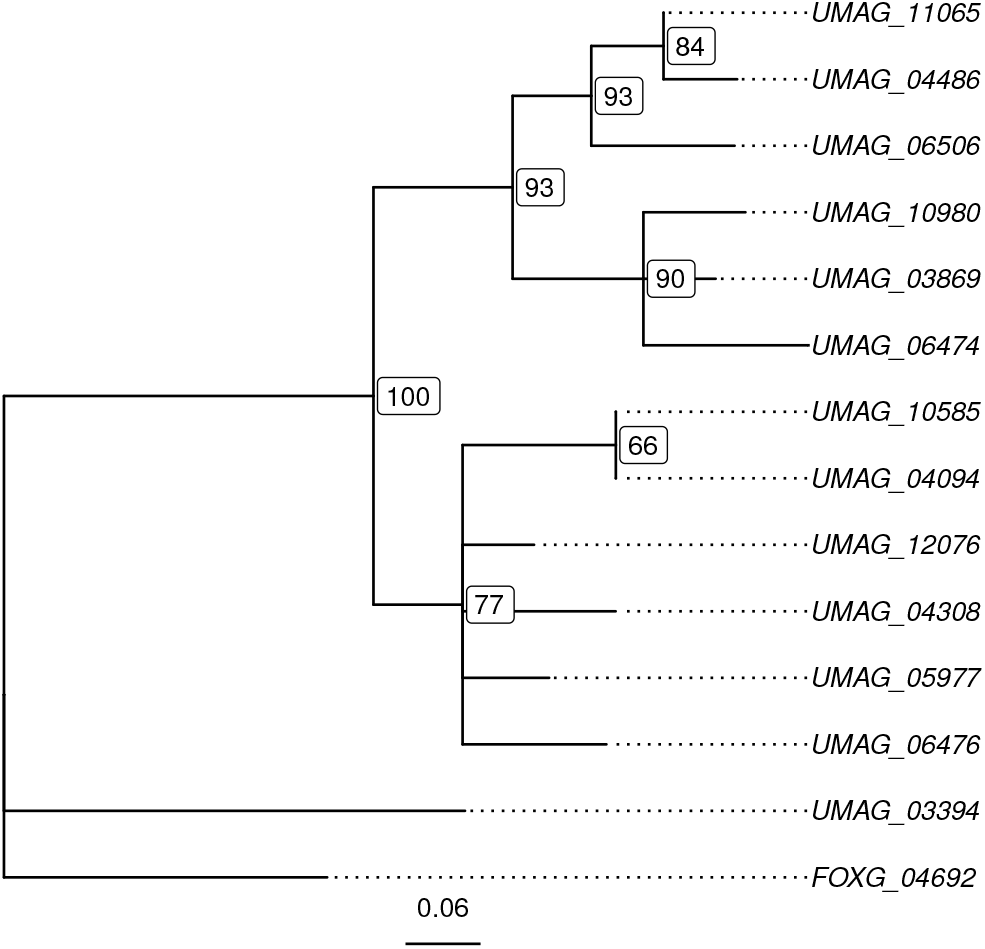
*Maximum likelihood phylogeny of* UMAG_11065 U. maydis *paralogs together with the closest homolog from* F. oxysporum *(see Table 1). Nodes labels indicate support values as percentage. Nodes with support values lower than 60% have been collapsed*.

### 5 *U. maydis* populations shows structural polymorphism in the telomeric region of chromosome 9

Because the *UMAG_11064* gene still displays a strong signature of its mitochondrial origin (codon usage and GC content), its transfer may have occurred recently. In order to provide a timeframe for the insertion event, we examined the structure of the genomic region of the insertion in other *U. maydis* and *S. reilianum* isolates, as well as the structure of the *cox1* exons 1, 2 and 7. The regions that could be amplified and their corresponding sizes are listed in Supplementary Figure S2 and the inferred genome organisations are summarized in Figure 6. The UMAG_11064 gene is present in the FB1-derived strain SG200, as well as in the Holliday strains 518 and 521, but is absent in the nuclear and mitochondrial genome sequences of a recent *U. maydis* isolate from the US, strain 10-1, as well as from 5 Mexican isolates (I2, O2, P2, S5 and T6, Figure S2A). Conversely, the *UMAG_11072* gene, which is located further away from the telomere on the same chromosome arm, could be amplified in all strains. This positive control demonstrates that the lack of amplification of *UMAG_11064* in some strains is not due to any issue with the quality of the extracted DNA (Figure S2B). These results suggest that, either the *UMAG_11064* gene was ancestral to all tested strains and subsequently lost in the Mexican and 10-1 strains, or it inserted in an ancestor of the three strains 518, 521 and SG200, after the divergence from other U. maydis strains, an event that occurred after the domestication of maize and the spread of the associated pathogen, 10,000 to 6,000 years ago (Munkacsi et al. 2008). Moreover, all U. maydis strains possess intron 6 in the mitochondrial cox1 gene, which is absent in S. reilianum. While the three S. reilianum strains tested carry intron 1, the most direct descendant of the progenitor of the HEs, it was absent in all U. maydis strains tested (Figure S2C-E and Figure 6).

**Figure 6.**
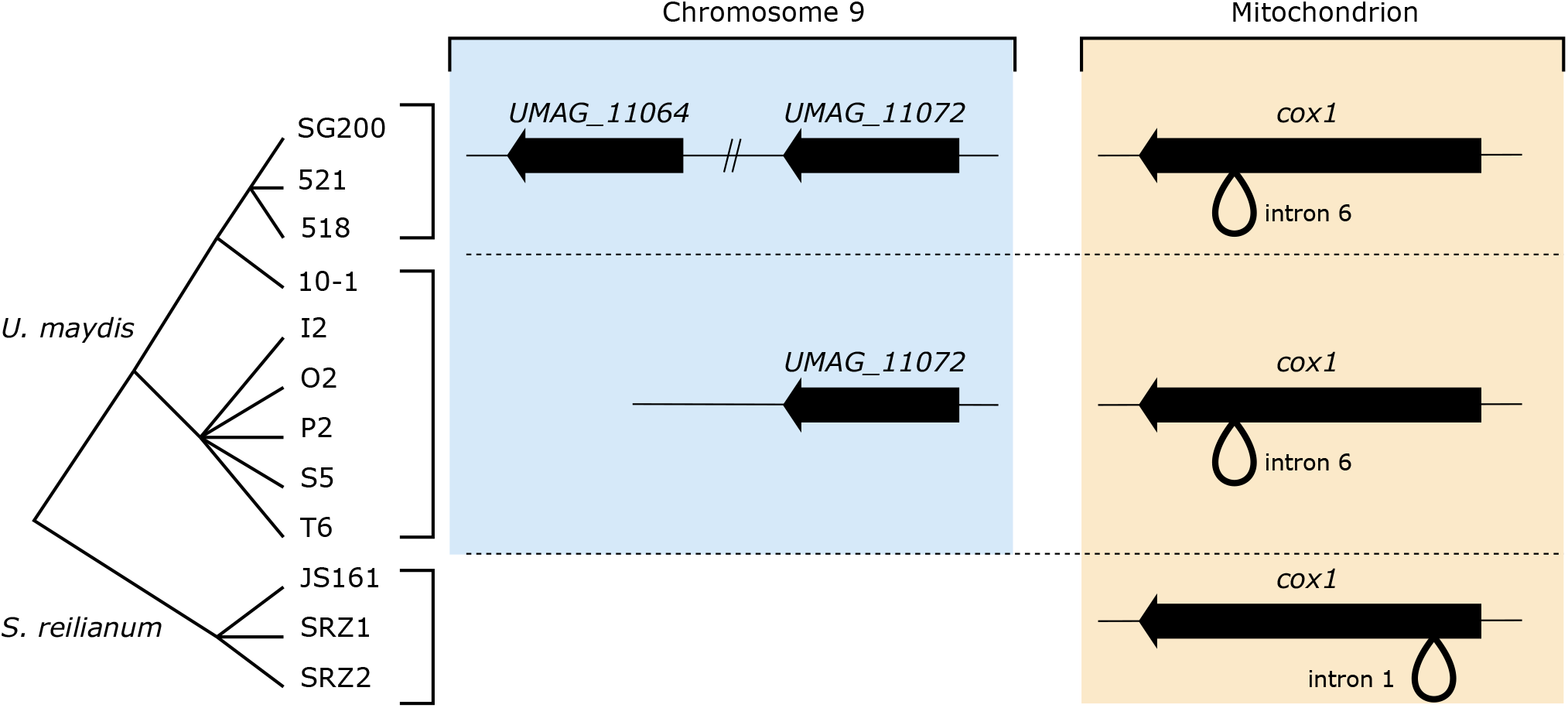
*Presence of the* UMAG_11064 *gene and structure of the* cox1 *gene in several* U. maydis *and* S. reilianum *strains, as assessed by PCR, together with their phylogeny. The* UMAG_11072 *gene, located 90 kb downstream the* UMAG_11064 *gene on chromosome 9, was used as a positive control. Strains 521 and 518 are two strains resulting from the same spore from a field isolate from the U.S.A. SG200 is a genetically engineered strain derived from a cross between the 518 and 521 strains. Strain 10-1 is another field isolate from the U.S.A. Strains I2, O2, P2, S5 and T6 from field isolates from Mexico*.

### 6 Functional characterization

To shed light on the functional implication of the translocation of the HEG and subsequent mutations we (i) assessed the expression profile of these genes and (ii) generated a deletion strain and phenotyped it. For the expression analysis we relied on a previously published RNASeq data set (Lanver et al. 2018), from which we extracted the expression profiles of genes in the telomeric region of chromosome 9 (Figure 7A). While the expression of *UMAG_11064* remained close to zero in the three replicates, expression of *UMAG_11065* increased during plant infection. The telomeric region was highly heterogeneous in terms of expression profile: while *UMAG_11066* and *UMAG_03393* did not show any significant level of expression, *UMAG_03392* was down-regulated at twelve hours post-infection, while *UMAG_03394*, another RecQ-encoding gene homologous to *UMAG_11065*, displayed constitutively high levels of expression (Figure 7A). All homologs of *UMAG_11065* show a significantly higher expression during infection (Tukey’s posthoc test, false discovery rate of 5%, Figure 7B). The comparison of expression profiles revealed two main classes of genes (Figure 7C): highly expressed genes (upper group), and moderately expressed genes (lower group), to which *UMAG_11065* belongs. We further note that the differences in expression profiles do not mirror the protein sequence similarity of the genes (Mantel permutation test, p-value = 0.566).

**Figure 7.**
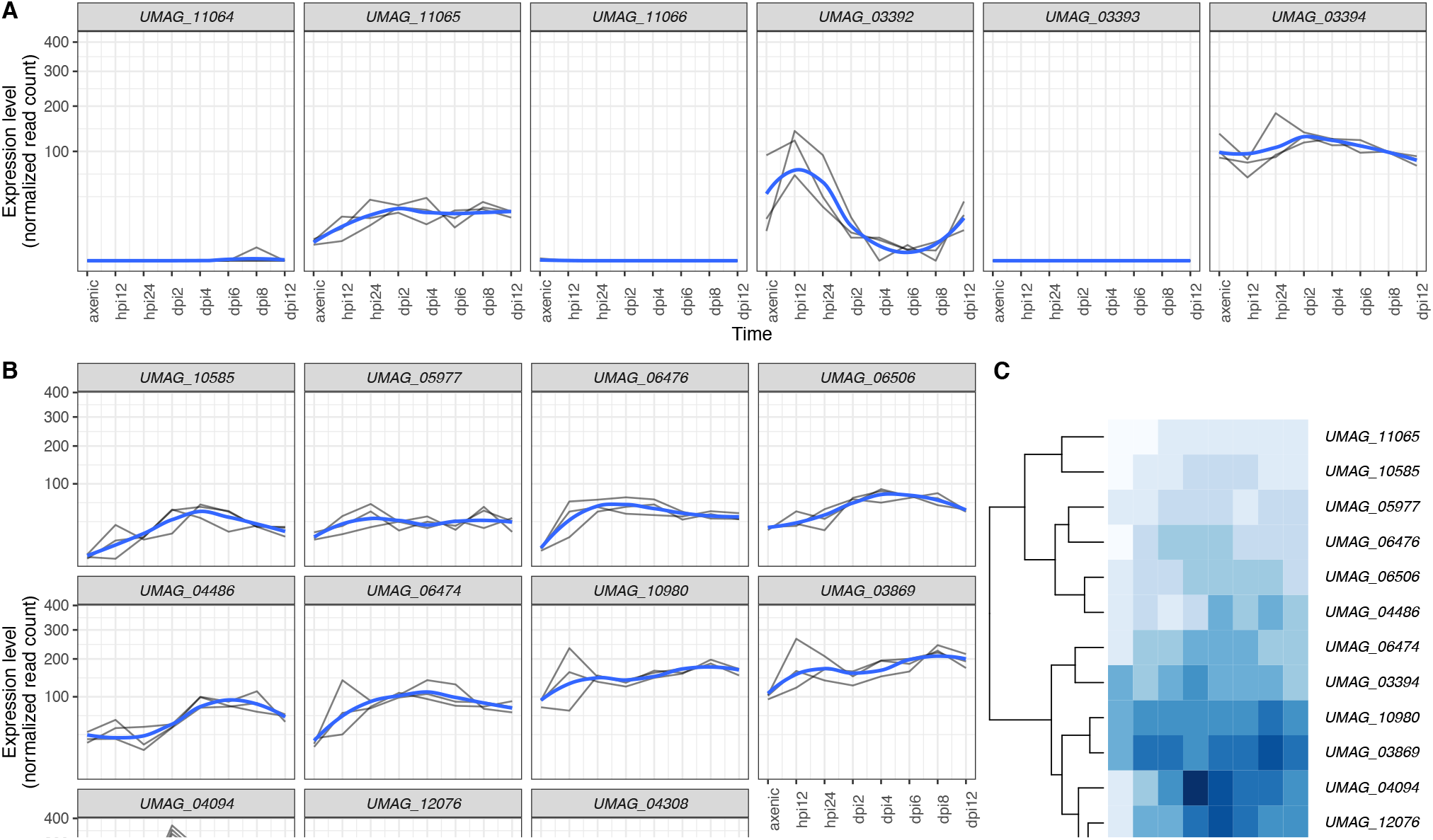
*Patterns of gene expression for* UMAG_11064 *and* UMAG_11065*, together with neighboring and homologous genes. A) Gene expression profiles for genes in the chromosome 9 telomeric region (as depicted on Figure 1B). Straight lines represent three independent replicates, while the blue curve depicts the smoothed conditional mean computed using the LOESS method. B) Gene expression profiles for the* UMAG_11065 *homologs (Figure 5). Legends as in A. C) Clustering of the* UMAG_11065 *homologs based on their averaged expression profile (see Methods). Hpi: hours post-infection. Dpi: days post-infection*.

To assess the functional role of *UMAG_11064* and *UMAG_11065*, these genes were simultaneously deleted in SG200, a solopathogenic haploid strain that can cause disease without a mating partner (Kämper et al. 2006) using a single-step gene replacement method (Kämper 2004). Gene deletion was verified by Southern analysis (Figure S3). Virulence assays, conducted in triplicate revealed no statistically different symptoms of the double deletion strain, SG200Δ11065Δ11064, compared to SG200 in infected maize plants (Figure 8A, Chi-square test, p-value = 0.453). Since RecQ helicases contribute to dealing with replication stress (Kojic and Holloman 2012) we also determined the sensitivity of the mutant to various stressors including UV, hydroxyurea and Congo Red. (Figure 8B). We report that the deletion strain shows increased sensitivity to cell wall stress induced by Congo Red and increased resistance to UV stress. Since *UMAG_11064* does not show any detectable level of expression, we hypothesize that the deletion of *UMAG_11065* is responsible for this phenotype.

**Figure 8.**
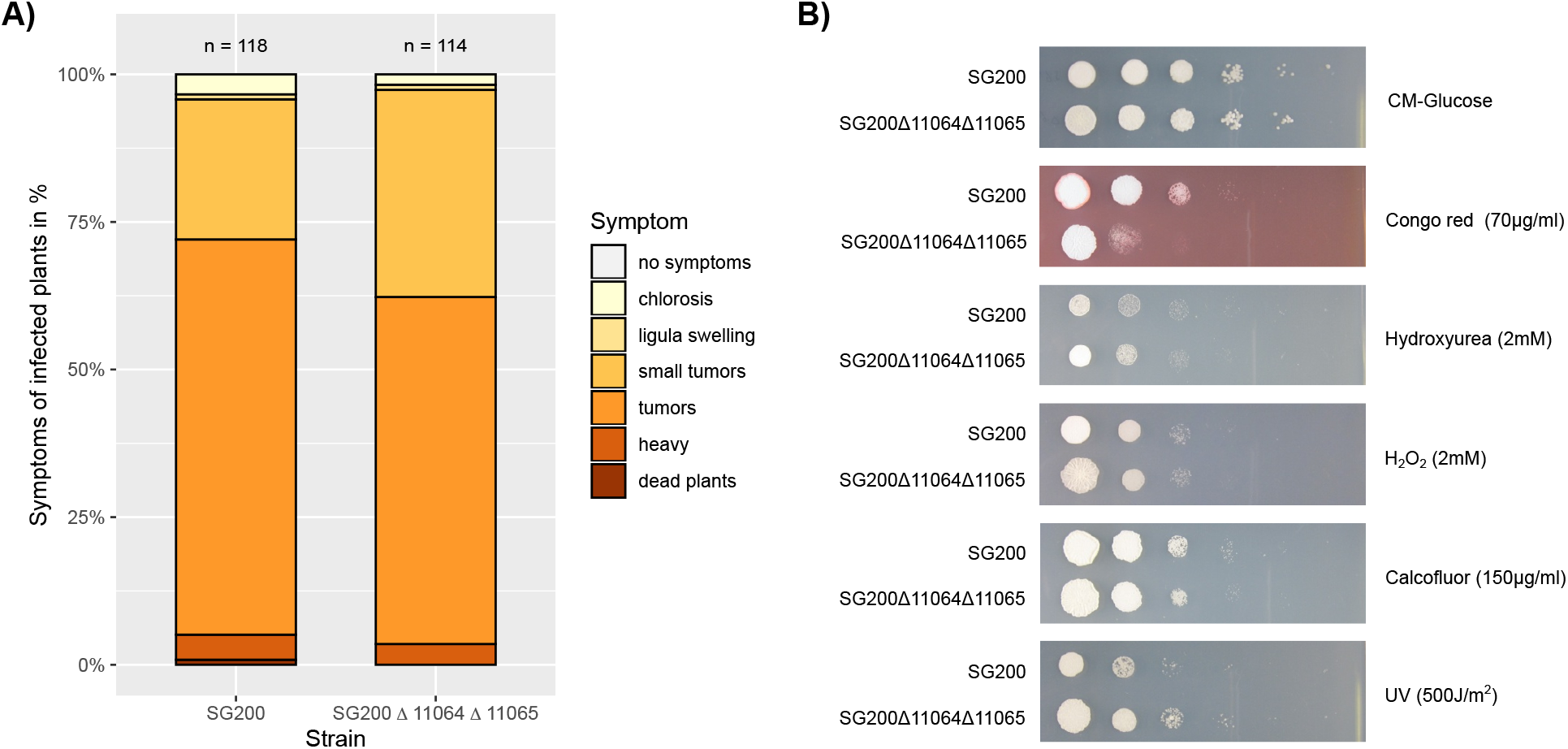
*Phenotype assessment of the double deletion strain. A) The simultaneous deletion of* UMAG_11064 *and* UMAG_11065 *does not affect virulence. Maize seedlings were infected with the indicated strains. Disease symptoms were scored at 12 dpi according to Kämper et al. (Kämper et al. 2006) using the color code depicted on the right. Colors reflect the degree of severity, from brown-red (severe) to light yellow (mild). Data represent mean of n = 3 biologically independent experiments. Total numbers of infected plants are indicated above the respective columns. B) Stress assay of the double deletion strain (Δ11064Δ11065), lacking both genes* UMAG_11064 *and* UMAG_11065, *compared to the parental SG200 strain. Assays were repeated at least three times with comparable results*.

## Discussion

The codon usage and GC content of the *UMAG_11064* gene, as well as its similarity to known mitochondrial HEGs, points at a recent transfer into the nuclear genome of *U. maydis*. Moreover, the precursor of this gene is absent from the mitochondrial genome of this species. Two possible scenarios can explain this pattern, which we detail below.

The first scenario involves a transfer of the gene to the nuclear genome followed by a loss of the mitochondrial copy (Figure 9). Under this scenario, the mitochondrial HEG was present in the *U. maydis* ancestor. Two evolutionary events are invoked: the insertion of the HEG into the nuclear genome, on the one hand, creating a HEG^+^ genotype at the nuclear locus (designated [HEG^+^]_nuc_), and the loss of the mitochondrial copy, creating a HEG^-^ genotype at the mitochondrial locus (designated [HEG^-^]_mit_). These two events may have happened at distinct time points, but, under this scenario, the former cannot have happened after the fixation of the [HEG^-^]_mit_ genotype in the population. The [HEG^+^]_nuc_ / [HEG^-^]_mit_ genotype could be generated by a cross between two individuals, one [HEG^+^]_nuc_ and the other [HEG^-^]_mit_, given that mitochondria are uniparentally inherited in *U. maydis* (Basse 2010). Importantly, the segregation of the [HEG^+^]_nuc_ and [HEG^-^]_mit_ variants could be purely neutral and driven by genetic drift only. In case the [HEG^-^]_nuc_ allele contained a recognition sequence of the HE, the [HEG^+^]_nuc_ allele may have initially benefited from a genetic drive effect. Any putative selective advantage/disadvantage of the [HEG^+^]_nuc_ or [HEG^-^]_mit_ alleles may have favored their fixation, or on the contrary, acted against it.

**Figure 9.**
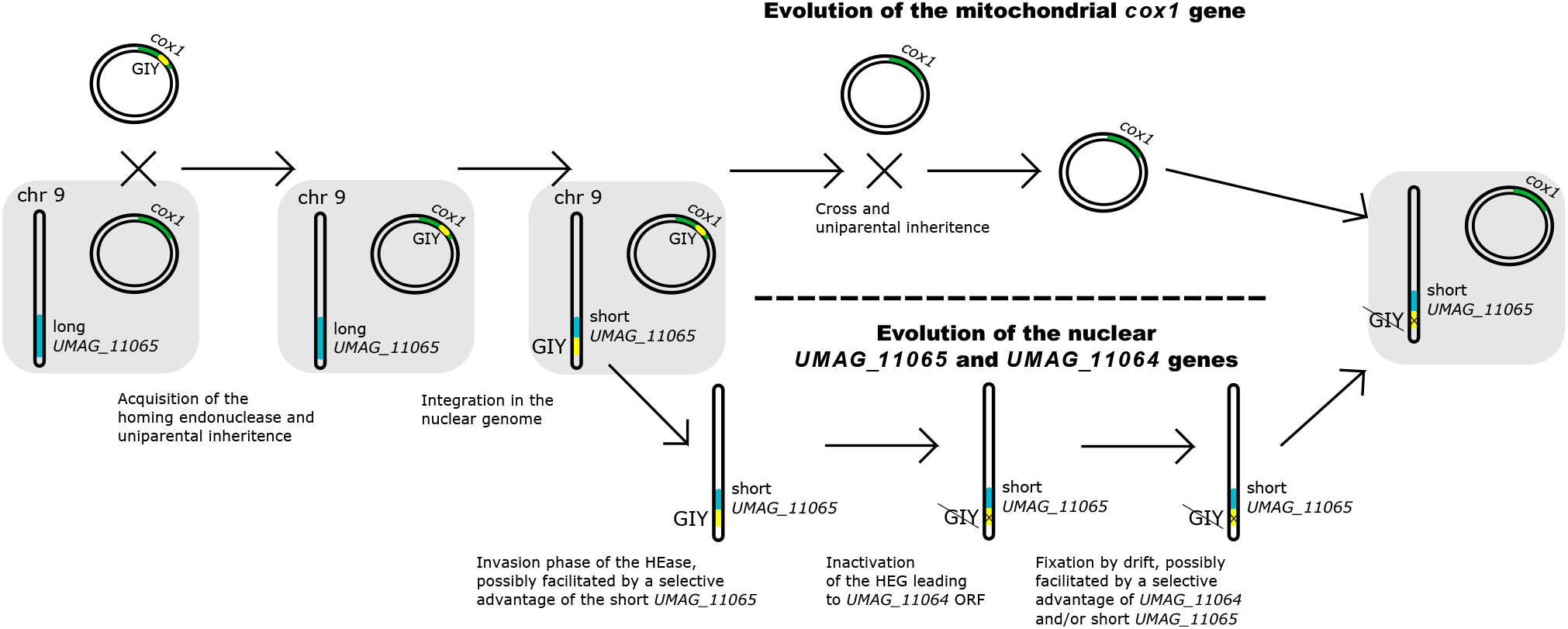
*Possible evolutionary scenario recapitulating the events leading to the formation of the* UMAG_11064 *and* UMAG_11065 U. maydis *genes. Importantly, each step in this model may have occurred by genetic drift alone. Positive selection may have favored - but is not required to explain - the spread of the nuclear and/or the loss of the mitochondrial HEGs*.

In the second scenario, the mitochondrial HEG was not ancestral to *U. maydis*, but was horizontally transferred from *S. reilianum* (or a related species). The high similarity of the *UMAG_11064* gene to the *S. reilianum* mitochondrial HEG (Figure 3) supports this hypothesis, given the relatively high nucleotide divergence between the two species, which diverged around 20 My ago (Schweizer et al. 2018). We note, however, that intronic HEGs have also been reported to show reduced nucleotide substitution rates, which can potentially explain their comparatively low divergence (Jalalzadeh et al. 2015). Group I introns have been reported to be highly mobile and to undergo frequent horizontal gene transfers (HGT) within metazoans (Schuster et al. 2017), plants (Sanchez-Puerta et al. 2008) and fungi (Jalalzadeh et al. 2015; Férandon et al. 2010). Group I introns in metazoans and plants are also thought to originate from a fungal donor (Schuster et al. 2017; Sanchez-Puerta et al. 2008, 2011). In this respect, a transfer from another *cox1* intron-carrying smut fungus to *U. maydis* is not unlikely, given that *U. maydis* and *S. reilianum* share the same host, and that hybridization between smut species has been reported (Fischer 1957; Boidin 1986).

The insertion of the HEG in the nuclear genome poses the question of the underlying mechanisms, independently of the origin of the HEG. First, the HEG could be encoding a fully functional HE and the [HEG^-^]_nuc_ allele contained a HE recognition sequence. Under this scenario, the insertion event was a homing event and the inactivation of the HEG occurred after the insertion. Therefore, several generations must have passed since the insertion event in order for the inactivating mutations to occur. Alternatively, the [HEG^-^]_nuc_ allele might not have contained a recognition sequence and the insertion of the HEG occurred by an unknown mechanism. Lastly, a possibility is that the inserted sequence already encoded an inactivated HE, which was then inserted by an unknown mechanism. In this latter case, the insertion could have occurred very recently, possibly a few generations in the past.

HEGs are found in eukaryotic nuclei but are usually restricted to small and large ribosomal RNA subunit genes (Lambowitz and Belfort 1993; Dunin-Horkawicz et al. 2006). While transfer of DNA segments and functional genes from organellar genomes to the nucleus is well documented (Sun and Callis 1993; Thorsness and Weber 1996; Lloyd and Timmis 2011; Fuentes et al. 2012), established examples of HEG insertions at other genomic locations than rRNA genes is very scarce. Several questions remain unanswered regarding the mechanisms of insertion into the nuclear genome, providing it happened as a homing event. For the event to happen, a template sequence containing the HEG is required for repairing the break initiated by the HE. This template must have, therefore, “leaked” from the mitochondrion. The recognition motif of the original *UMAG_11064* HEG is unknown, and the very short flanking regions surrounding the insertion site do not allow any comparison with known motifs, preventing further conclusions to be made regarding the nature of the insertion event of *UMAG_11064*.

Interestingly, Louis and Haber (Louis and Haber 1991) reported a similar transfer of a HEG into a telomeric region of *Saccharomyces cerevisiae*. The authors argue that signatures of such insertion could be found because (i) it had no deleterious effect and (ii) the occurrence of heterologous recombination between telomeres favours the maintenance of elements that would otherwise be lost. Contrasting with this result, the insertion of the GIY-YIG HEG that inserted into the ancestor of the *UMAG_11065* gene potentially had nonneutral effects, resulting in an expressed truncated protein. Several mutations were found within the active site of the inserted HEG that led to the *UMAG_11064* gene, suggesting that the encoded protein is unlikely to act as a HE any longer. However, a putative alternative start codon was detected, downstream the active site, followed by an uninterrupted peptide sequence containing the helix-turn-helix binding domain of the original HE. Furthermore, we could not detect any significant level of expression of the *UMAG_11064* gene in various laboratory conditions. Comparative sequence analysis further suggests that *UMAG_11064* is evolving under relaxed purifying selection, indicating that it might be undergoing pseudogenization. These results, therefore, suggest that the *UMAG_11064* gene is not functional. However, as this mitochondrial HEG inserted into a nuclear *U. maydis* gene, it might have had phenotypic consequences not directly due to the HEG gene itself. The *UMAG_11065* gene appeared truncated by the HEG insertion, which removed the C-terminal part of the encoded protein, a likely RecQ helicase, and the truncated *UMAG_11065* is expressed during infection. The truncation likely did not have a strong negative impact, possibly because of the existence of multiple potentially functionally redundant paralogs of *UMAG_11065*, including on the same telomeric region of chromosome 9, with *UMAG_03394* being located 4 genes upstream (Table 1). While we were unable to detect a contribution to virulence, our results point at a putative role of the truncated RecQ helicase into stress tolerance, as its deletion increases resistance to UV radiation but makes the fungus more susceptible to cell wall stress, at least under laboratory conditions. How the truncated UMAG_11065 RecQ helicase could improve coping with cell wall stress and increases the sensitivity to UV simultaneously, however, remains to be investigated, as well as the potential fitness benefit or cost of these phenotypes. Furthermore, a possibility remains that the observed phenotype of *UMAG_11065* is ancestral and not due to the truncation itself, which could be neutral. In order to elucidate the putative adaptive role of the truncation of *UMAG_11065*, knowledge of the ancestral, non-truncated *UMAG_11065* allele is needed, as well as its distribution in natural populations.

## Conclusions

In this study, we report instances of two stages of the life cycle of HEGs. Intron 1 of the mitochondrial *cox1* gene of *S. reilianum* was shown to contain a degenerated GIY-YIG HEG, while the homologous position in the *U. maydis* gene displays no intron. Besides, in the telomeric region of chromosome 9 of the nuclear genome of *U. maydis*, we found evidence of a recent insertion of a very similar GIY-YIG HEG. Phenotypic assay of the mutant strain containing a double-deletion of the HEG and the helicase gene where it inserted reveals enhanced stress sensitivity *in vitro*. The absence of a GIY-YIG HEG in any field isolates of *U. maydis* sequenced so far, however, suggests that either the mutation was lost in natural populations and only maintained under laboratory conditions, or that it is only present in a so far unsampled population. The *UMAG_11064* gene offers a snapshot of evolution taken soon after a mutation event occurred. As such, it can provide insights into the mechanisms of HEG mobility and horizontal transfer. These results further demonstrate that HEGs can generate genetic diversity not only via their duplication, but also by drastically modifying the local genome architecture where they insert.

## Data accessibility

Data sets and scripts necessary to reproduce the statistical analyses in this work are available as Supplementary File S1 and at https://gitlab.gwdg.de/molsysevol/umag_11064.

## Supplementary material

**Supplementary Table S1:** Homology search results using *UMAG_11064* as a query on the NCBI non-redundant nucleotide database, using BlastN. All hits with an E-value lower than 1E-04 are included, alongside with corresponding alignment length and percentage of sequence identity.

**Supplementary Table S2:** Homology search results using *UMAG_11064* as a query on NCBI non-redundant protein database, using BlastP. All hits with an E-value lower than 1E-04 are included, alongside with corresponding alignment length and percentage of sequence identity.

**Supplementary Table S3:** Homology search results using *UMAG_11065* as a query on NCBI non-redundant protein database, using BlastP. All hits with an E-value lower than 1E-04 are included, alongside with corresponding alignment length and percentage of sequence identity.

**Supplementary Table S4:** Homology search results using *U. maydis cox1* intron 6 as a query on a NCBI non-redundant protein database, using BlastX. All hits with an E-value lower than 1E-04 are included, alongside with corresponding alignment length and percentage of sequence identity.

**Supplementary Table S5:** Homology search results using *S. reilianum cox1* intron 1 as a query on a NCBI non-redundant protein database, using BlastX. All hits with an E-value lower than 1E-04 are included, alongside with corresponding alignment length and percentage of sequence identity.

**Supplementary Table S6:** Homology search results using *S. reilianum cox1* intron 2 as a query on a NCBI non-redundant protein database, using BlastX. All hits with an E-value lower than 1E-04 are included, alongside with corresponding alignment length and percentage of sequence identity.

**Supplementary Table S7:** Primers used in this study.

**Supplementary Figure S1:** Amplification of the candidate region in the telomeric region of chromosome 9. A) Genomic context based on the *U. maydis* reference genome, and location PCR primers. B) PCR results with corresponding expected fragment sizes. Forward (fw) and reverse (rv) primer sequences are provided in Table S7.

**Supplementary Figure S2:** Amplification of A) *UMAG_11064*, B) *UMAG_11072* and *cox1* exons C) 1 and D) 7 in several *U. maydis* and *S. reilianum* strains. Forward (fw) and reverse (rv) primer sequences are provided in Table S7. E) Summary table of the results. Plus and minus signs indicate whether the corresponding gene could be amplified or not. Numbers indicate the size of the amplified region in base pairs. Strains are labelled as in Figure 6.

**Supplementary Figure S3**: Verification of the deletion of *UMAG_11064* and *UMAG_11065*. A) Schematic map of the genomic region containing *UMAG_11064* and *UMAG_11065* in SG200 and SG200D11064D11065. Primers used to amplify the left and right border sequences are indicated. B) DNA of SG200 and SG200Δ11064Δ11065 was cleaved with Fsp1 and subjected tho southern blot analysis using a mixture of Probes 1 and 2 indicated in A). The 2.94 kb fragment is diagnostic for SG200 while the 4.19 kb fragment is diagnostic for the deletion of *UMAG_11064* and *UMAG_11065*.

**Supplementary File S1**: Scripts used to conduct the phylogenetic and statistical analyses, As well as R code used to generate figures 1, 3, 4, 5 and 6.

## Acknowledgements

We thank all members of our groups for stimulating discussions. We are grateful to Georgiana May and Octavio Paredes-López for providing field isolates of *U. maydis* from the US and Mexico, respectively. We acknowledge the generous support by the Max Planck Society. Version 4 of this preprint has been peer-reviewed and recommended by Peer Community In Evolutionary Biology (https://doi.org/10.24072/pci.evolbiol.100101)

