## Supplementary figures and images for "The insertion of a mitochondrial selfish element into the nuclear genome and its consequences"

### MSA-PhyML_tree.pdf

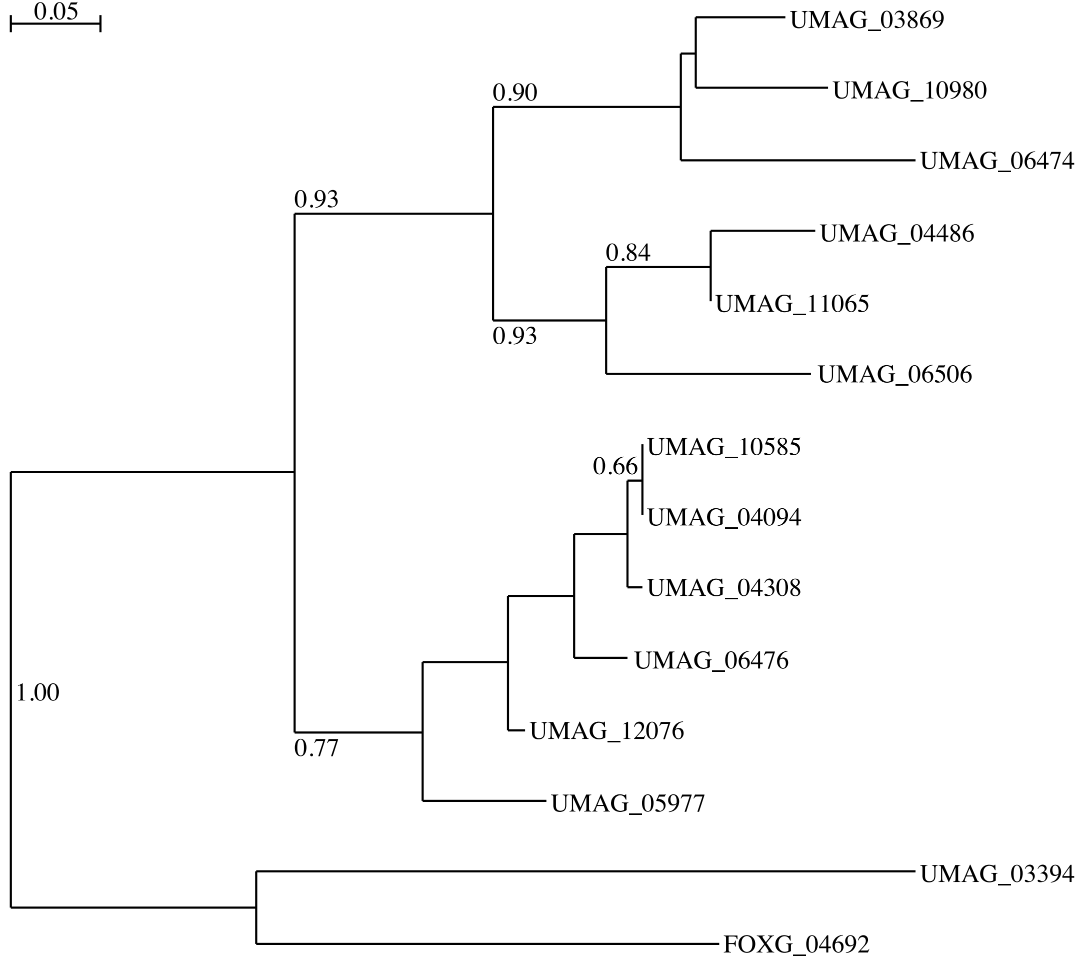

### Supplementary Figure S1

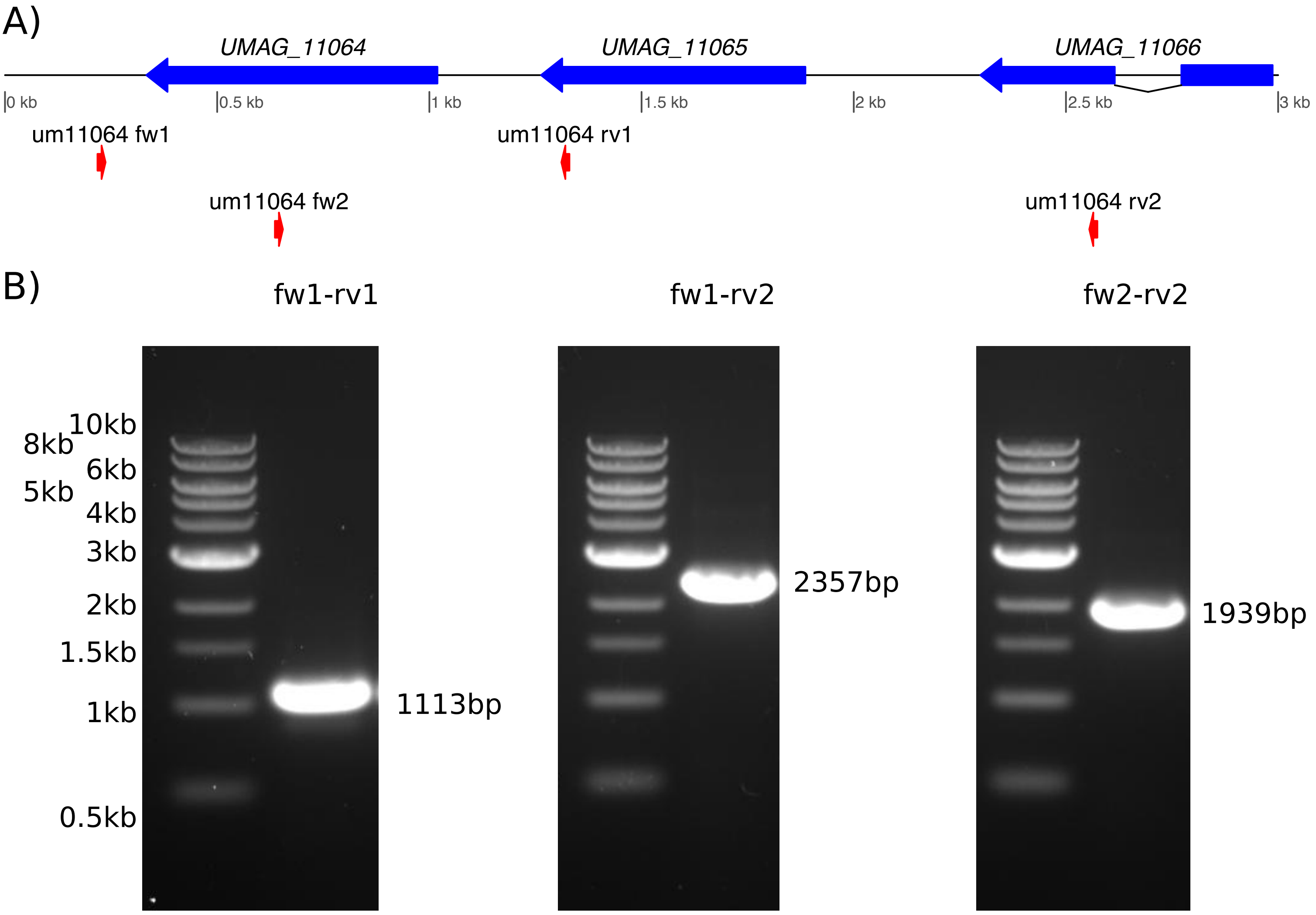

### Supplementary Figure S3

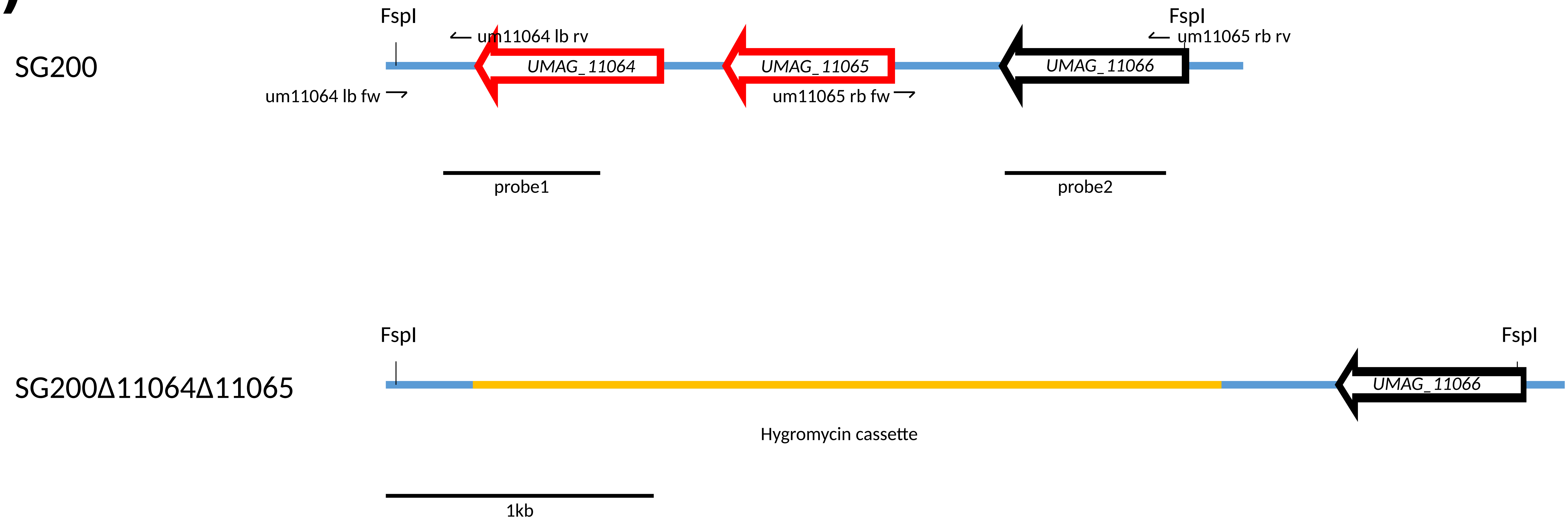

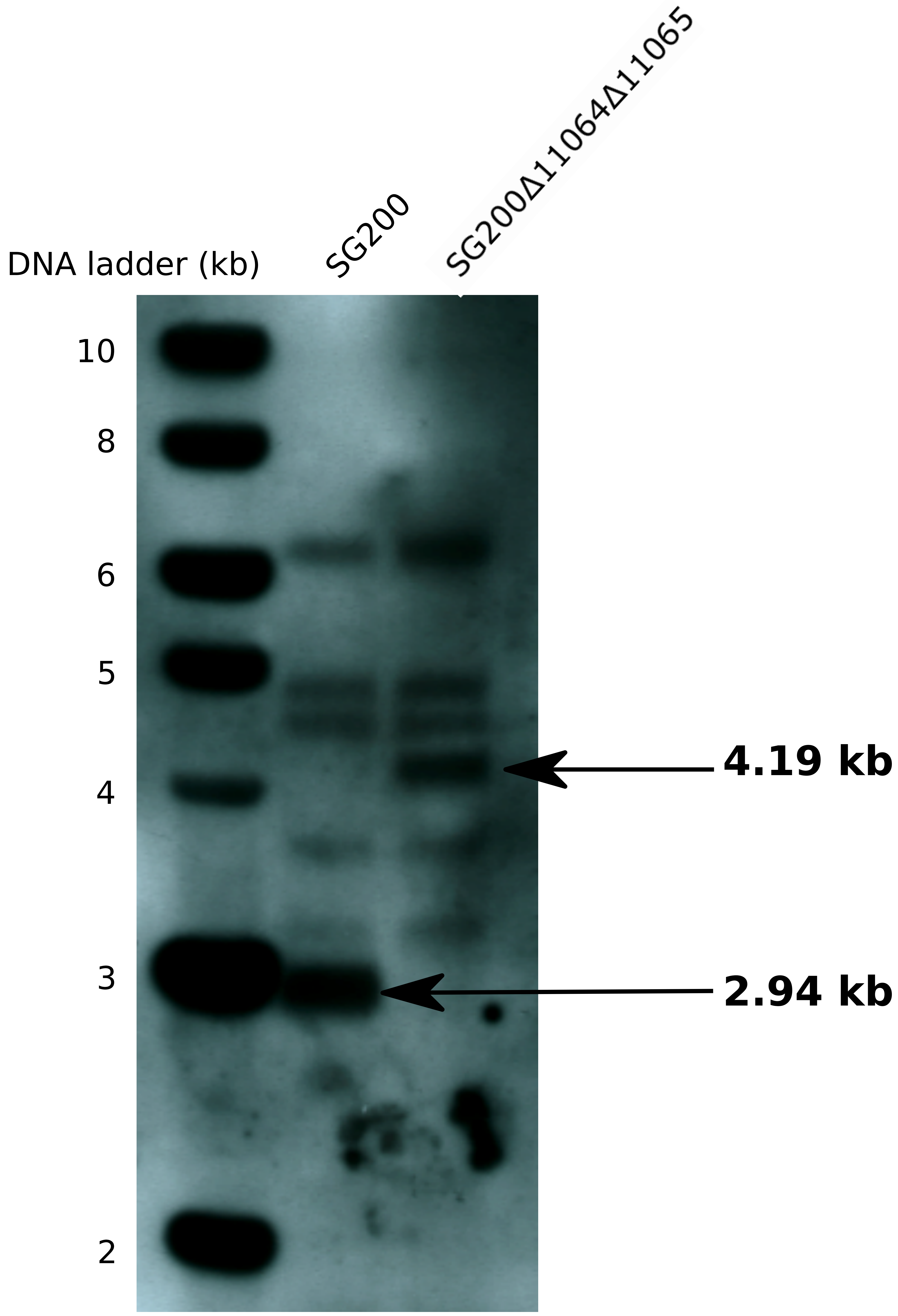
