## Supplementary Figure S2 for "The insertion of a mitochondrial selfish element into the nuclear genome and its consequences"

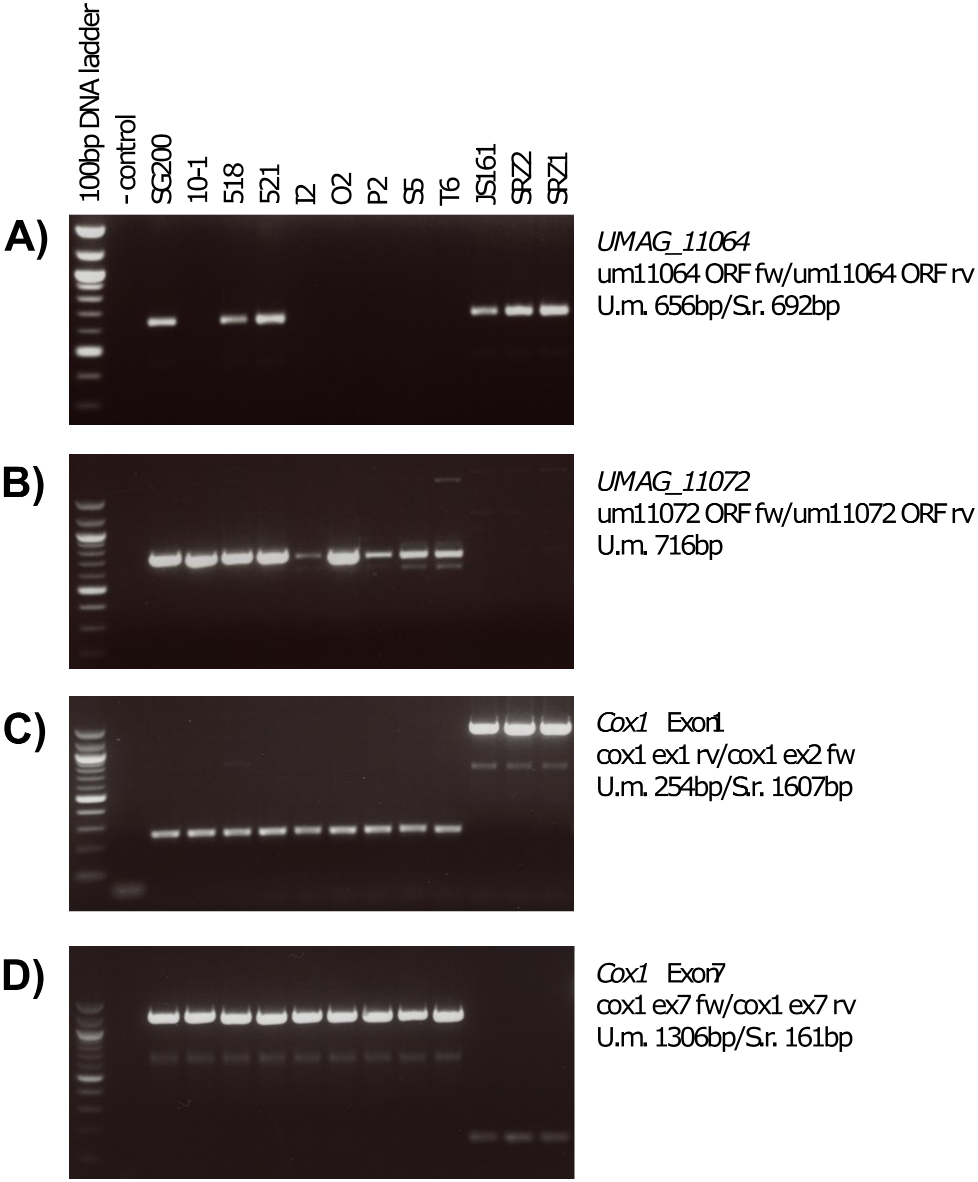


**E)**

| Region | Strain | | | | | | | | | | | |
|---|---|---|---|---|---|---|---|---|---|---|---|---|
| | *U. maydis* | | | | | | | | | *S. reilianum* | | |
| | SG200 | 10-1 | 518 | 521 | I2 | O2 | P2 | S5 | T6 | JS161 | SRZ1 | SRZ2 |
| *UMAG_11064* ORF | + | - | + | + | - | - | - | - | - | + | + | + |
| *UMAG_11072* ORF | + | + | + | + | + | + | + | + | + | (not tested) | | |
| *Cox1* Exon 1+2 | 254 | 254 | 254 | 254 | 254 | 254 | 254 | 254 | 254 | 1607 | 1607 | 1607 |
| *Cox1* Exon 7 | 1306 | 1306 | 1306 | 1306 | 1306 | 1306 | 1306 | 1306 | 1306 | 161 | 161 | 161 |
