## Supplementary File S1 for "The insertion of a mitochondrial selfish element into the nuclear genome and its consequences": parseBlast.py

```
#! /usr/bin/python

from Bio.Blast import NCBIXML

#result_handle = open("J4K2HP2M015-Alignment.xml") #no model/no uncultured
result_handle = open("JBJS4UZ501N-Alignment.xml")
blast_records = NCBIXML.parse(result_handle)

print("Name,Description,AlignmentLength,PercentIdentity,HSPexpect")
for blast_record in blast_records:
  for alignment in blast_record.alignments:
    for hsp in alignment.hsps:
      if hsp.expect < 0.0001:
        t = alignment.title.split(">")[0].split(" ")
        name = t[0]
        name = name.split("|")[3]
        t.pop(0)
        desc = " ".join(t)
        print "%s,'%s',%d,%f,%s" %(name, desc, hsp.align_length, float(hsp.identities) / float(hsp.align_length), hsp.expect)
```
