## Supplementary File S1 for "The insertion of a mitochondrial selfish element into the nuclear genome and its consequences": README.md

Created on 06/01/20 by jdutheil

1. Get sequences
================

Retrieved sequences from NCBI (IDs are provided in Table S2).
Add nucleotide sequence for Sr, Ti, Tw, Ss and Ub (Table S1). Complement in Seaview (Sr only).
Rename and remove duplicated sequence from A. bisporus (original sequence name in `_orig` file).

2. Alignment using Macse
========================

```bash
java -Xmx600m -jar ~/.local/bin/macse_v1.2.jar -prog alignSequences -gc_def 4 -seq UMAG_11064relatives.fasta
```

3. Phylogeny reconstruction
===========================

Selection done in Seaview, using manual editing.
Phylogeny using PhyML in Seaview, Gamma(4) model, best of NNI and SPR.
The GTR model was used for nucleotide sequence (all species), and the LG model for protein sequences (after excluding the obvious pseudogenes). For the nucleotide sequences, ! were replaced by - in the fasta alignment (*selection_allSpecies.fasta). 
100 non-parametric boostrap. Bio++ PhyView was then used to collapse node with support lower than 65% in the protein alignment, and to further remove remaining bootstrap values.

4. Positive selection analysis
==============================

Convert sequence to PAML format:
```bash
bppseqman param=SeqMan.bpp DATA=UMAG_11064relatives_macse_NT_selection
bppseqman param=SeqMan.bpp DATA=UMAG_11064relatives_macse_NT_selection2
```

Run PAML:
```bash
codeml
```
There seems to have convergence issues... We run it 10 times and check:
```bash
for i in {1..10}; do
  echo "Estimation $i..."
  codeml
  cp UMAG_11064_branch.mlc UMAG_11064_branch_$i.mlc
done;
```

Check the results:
```r
require(treeio)
require(data.table)
dat.lst <- list()
for (i in 1:10) {
  mlc <- read.codeml_mlc(paste("UMAG_11064_branch_", i, ".mlc", sep = ""))
  dat <- mlc@data
  dat$Rep <- i
  dat.lst[[i]] <- dat
}
dat <- rbindlist(dat.lst)

require(plyr)
ddply(.data = dat, .variables = "node", summarize, Min = min(dN_vs_dS), Max = max(dN_vs_dS))

plot(dN_vs_dS~node, dat, log = "y"); abline(h=1)
plot(dN~node, dat, log = "y"); abline(h=1)
plot(dS~node, dat, log = "y"); abline(h=1)
```

We try with bppML:
```bash
bppml param=BppML.bpp
```

For checking... (only 1 foreground sequence)
Use a branch site model:
```bash
codeml codeml2.ctl
codeml codeml2null.ctl
```
Run multiple times: no convergence issue here, good.

```r
l0 <- -4882.500811
l1 <- -4881.506731

s <- 2 * (l1 - l0)
pchisq(s, df = 1, lower.tail = FALSE)
```

pvalue: 16% NS.


We remove the two Tilletia from the background (selection 2):
```bash
codeml codeml2b.ctl
codeml codeml2bnull.ctl
```

```r
l0 <- -4235.880736
l1 <- -4235.122847

s <- 2 * (l1 - l0)
pchisq(s, df = 1, lower.tail = FALSE)
```

pvalue: 22%

5. Make a figure
================

```r
require(ggtree)
tree <- read.tree("UMAG_11064relatives_macse_NT_selection-PhyML_tree_allspecies.dnd")
p <- ggplot(tree, aes(x, y))
p <- p + geom_tree() + geom_tiplab() + geom_nodelab(geom = "label", hjust = 0) + geom_treescale()
p <- p + theme_tree() + xlim(0,1)
p
ggsave(p, filename = "Phylogeny.pdf", width = 8, height = 6)
```
The rest we edit in inkscape.
