## Supplementary File S1 for "The insertion of a mitochondrial selfish element into the nuclear genome and its consequences": AnalyzeUsageUstilagoMaydis.R

```
# Created on 23/04/12 by jdutheil
# Analyze of codon usage in Ustilago maydis
# Modified from http://pbil.univ-lyon1.fr/datasets/charif04/

####################################################################
#
# Load required packages, here ade4 for multivariate analyses and seqinR
# to access ACNUC databases. This is dispensable since these packages
# are automatically pre-loaded in our RWeb server, but would be mandatory
# for runing the script in a different context.
#
####################################################################

library(ade4)
library(seqinr)

####################################################################
#
# Give session info to check that we are running the last version
# of R and the packages.
#
####################################################################

sessionInfo()

####################################################################
#
# Get sequences:
#
####################################################################

seq.um <- read.fasta("Umaydis_valid_orf_corrected.fa", seqtype="DNA")
seq.um.mito <- read.fasta("NCBI_mitochondrion_orf.fasta", seqtype="DNA")

####################################################################
#
# From sequences, build dataframes with codon usage for each sequence. 
#
####################################################################

mkdata <- function(seqs)
{
  tab <- sapply(seqs, uco)  
  tab <- as.data.frame(tab)
  return( tab )
}

tab.um <- mkdata(seq.um)
tab.um.mito <- mkdata(seq.um.mito)

####################################################################
#
# Remove CDS with in-frame stop codons (translated by "*" here)
# and CDS that have less than 100 codons:
#
####################################################################

cleanup <- function(tab)
{
  aa <- translate(sapply(rownames(tab),s2c))
  tab <- tab[ , colSums(tab[which(aa == "*"), ]) == 1]
  tab <- tab[ , colSums(tab) > 100 ]
  cat(ncol(tab), " cds.\n")
  return(tab)
}

tab.um  <- cleanup(tab.um)  # now 6,595 cds
tab.um.mito <- cleanup(tab.um.mito) # now 22 cds

####################################################################
#
# Run correspondence analysis on merged dataset:
#
####################################################################

tab <- cbind(tab.um, tab.um.mito)
#names(tab) <- 1:ncol(tab)
coa <- dudi.coa(tab, scan = FALSE, nf = 4)

####################################################################
#
# Run synonymous codon usage analysis:
#
####################################################################

facaa <- as.factor(aaa(translate(sapply(rownames(tab),s2c))))
scua <- wca(coa, facaa, scan = FALSE, nf = 4)

####################################################################
#
# Plot first factorial map:
#
####################################################################

l<-matrix(nrow=2, ncol=2)
l[1,]<-1
l[2,1]<-2
l[2,2]<-3
layout(l, widths=c(4,1))

###
# Figure A
#

par(mar=c(4,4,1,0)+0.1)
pch=c(Nuclear=4, Mito=18)
facsp <- as.factor(rep(c("Nuclear","Mito"), c(ncol(tab.um),ncol(tab.um.mito))))

plot(Comp2~Comp1, scua$co, xlab="First axis", ylab="Second axis", pch=pch[as.character(facsp)], xlim=c(-1, 1), main=NA)
title(main="A", adj=0) #For compound figure only
grid()
abline(h=0)
abline(v=0)
text(0.9, -0.7, "UMAG_11064", font = 3)
legend( x = -1, y = -0.5, pch = c(4, 18), col = "black", 
        legend = c(expression(italic("Nuclear")),
                   expression(italic("Mitochondrial"))), cex = 1, bty="n", xjust=0, y.intersp=1)

#dev.print(pdf, file="UmCodonUsage.pdf", width=10, height=6)

####################################################################
#
# END
#
####################################################################

# Analysis of GC content:

#l <- matrix(nrow = 1, ncol = 2)
#l[1,1] <- 1
#l[1,2] <- 2
#layout(l, widths = c(4,1))


seqs<-read.fasta("um.contig")

scangc<-function(chr, wsize, wstep, threshold=1) {
  res<-matrix(ncol=4, nrow=0)
  gc<-numeric(0)
  x<-numeric(0)
  b<-numeric(0)
  e<-numeric(0)
  pos<-0
  pb<-txtProgressBar(min=0, max=length(chr) - wsize, style = 3)
  while(pos + wsize < length(chr)) {
    setTxtProgressBar(pb, pos)
    val<-GC(chr[pos:(pos+wsize)])
    if (val <= threshold) {
      res<-rbind(res, c(val, pos + wsize/2, pos, pos + wsize))
    }
    pos<-pos + wstep
  }
  close(pb)
  res<-as.data.frame(res)
  names(res)<-c("gc", "pos", "start", "stop")
  return(res)
}

plot.gc<-function(gc, start, end, df, ...) {
  y<-gc$gc[gc$pos >= start & gc$pos < end]
  x<-gc$pos[gc$pos >= start & gc$pos < end]
  plot(y~x, type="l", col=grey(0.8), ...)
  lines(smooth.spline(x, y, df=df), col="black", lwd=2, type="l")
}

plot.gene<-function(name, start, end, col, y=0.65, w=0.05, tip=50, offset = 0.5) {
  if (all(end > start)) {
    cat("Gene in on the + strand.\n")
    endf<-max(end)
    end[length(end)]<-endf - tip
    rect(start, y, end, y+w, col=col, border=col)
    polygon(x=c(endf - tip, endf - tip, endf), y=c(y, y+w, y+w/2), col=col, border=col)
    end[length(end)]<-endf
  } else if (all(end < start)) {
    cat("Gene in on the - strand.\n")
    endf<-min(end)
    end[length(end)]<-endf + tip
    rect(start, y, end, y+w, col=col, border=col)
    polygon(x=c(endf + tip, endf + tip, endf), y=c(y, y+w, y+w/2), col=col, border=col)
    end[length(end)]<-endf
  } else {
    stop("Weird exon structure!!!")
  }
  text((max(end) + min(start)) / 2, y + w * offset, name, font = 3)
  segments(x0 = c(start, end),
           x1 = c(start, end),
           y0 = c(0, 0),
           y1 = c(y + w, y + w),
           lty = "dotted",
           col = gray(0.6))
}

gc.chr9<-scangc(seqs[["chr09"]][1:10000], 300, 1)
mito<-read.fasta("NCBI_mitochondrion_dna.fasta")
gc.mit<-scangc(mito[[1]], 300, 300)


par(mar=c(4.1,4.1,1.1,0))
plot.gc(gc.chr9, 0, 9000, df = 200, ylim = c(0.1,0.75), main = NA, xlab = "Chromosome position", ylab="% GC")
title(main="B", adj=0) #For compound image only
# Add gene annotations:
plot.gene("UMAG_11064", 1019, 333, grey(0.7), offset = -0.5)
plot.gene("UMAG_11065", 1886, 1263, grey(0.7), offset = 1.5)
plot.gene("UMAG_11066", c(2987, 2615), c(2772, 2298), grey(0.7), offset = 1.5)
plot.gene("UMAG_03392", 4106, 3495, grey(0.7), offset = 1.5)
plot.gene("UMAG_03394", 8836, 5960, grey(0.7), offset = 1.5)
plot.gene("UMAG_03393", c(5713, 5576, 5430), c(5623, 5463, 5222), grey(0.7), offset = 1.5)

abline(h=median(gc.mit$gc), lty="dashed")

#Finally compare GC distribution in nuclear vs. mitochondrial:
#gc.all<-scangc(seqs[["chr01"]], 1000, 1000)
par(mar=c(4.1,0,1.1,0.1))
boxplot(gc.mit$gc, ylim=c(0.1, 0.75), main=NA, xlab="Mitochondrial", yaxt="n")
#title(main="C", adj=0) #for compound figure only
#title(main="Mitochondrial GC", adj=0) #otherwise
abline(h=median(gc.mit$gc), lty="dashed")

#Single figure:
#dev.print(pdf, file="UmGCContent.pdf", width=10, height=6)

#Compound figure:
dev.print(tiff, file="CodonUsageAndGCContent.tif", width=3500, height=2500, compression="lzw", res=300)
```
