## Supplementary File S1 for "The insertion of a mitochondrial selfish element into the nuclear genome and its consequences": Analysis_GeneExpression.R

```
# Created on 04/07/19 by jdutheil
# Look at gene expression of UMAG_11064 and UMAG_11065

library(reshape2)
library(ggplot2)
library(plyr)
library(RColorBrewer)

dat <- read.table("GSE103876_NormCounts_Um.txt.gz", header = TRUE, stringsAsFactors = FALSE)

# First compare genes in the region:
dat2 <- subset(dat, Gene %in% c("UMAG_11064", "UMAG_11065", "UMAG_11066", "UMAG_03392", "UMAG_03393", "UMAG_03394"))
dat2$Gene <- ordered(dat2$Gene, levels = c("UMAG_11064", "UMAG_11065", "UMAG_11066", "UMAG_03392", "UMAG_03393", "UMAG_03394"))
dat3 <- melt(dat2, id.vars = "Gene")
dat3$replicate <- sapply(strsplit(as.character(dat3$variable), split = "_"), function(x) x[2])
dat3$time <- sapply(strsplit(as.character(dat3$variable), split = "_"), function(x) x[1])
dat3$time <- ordered(dat3$time, levels = c("axenic", "hpi12", "hpi24", "dpi2", "dpi4", "dpi6", "dpi8", "dpi12"))

p1 <- ggplot(dat3, aes(x=as.numeric(time), y=value, group = paste(replicate, Gene))) +
  geom_line(alpha = 0.5) + geom_smooth(se = FALSE, aes(group = Gene)) +
  scale_x_continuous(breaks = as.numeric(unique(dat3$time)), labels = levels(dat3$time)) +
  scale_y_sqrt(limits = c(0, 400)) +
  facet_grid(~Gene) +
  ylab("Expression level\n(normalized read count)") +
  xlab("Time") +
  theme_bw() +
  theme(strip.text = element_text(face = "italic"),
        axis.text.x = element_text(angle = 90))
p1

dat4 <- subset(dat, Gene %in% c("UMAG_11065", "UMAG_04486", "UMAG_06506", "UMAG_03394", "UMAG_03869", "UMAG_10980", "UMAG_06474", "UMAG_10585", "UMAG_04094", "UMAG_04308", "UMAG_06476", "UMAG_12076", "UMAG_05977"))
dat4$Gene <- ordered(dat4$Gene, levels = c("UMAG_11065", "UMAG_04486", "UMAG_06506",
                                           "UMAG_03869", "UMAG_10980", "UMAG_06474",
                                           "UMAG_05977", "UMAG_12076", "UMAG_06476",
                                           "UMAG_04308", "UMAG_04094", "UMAG_10585",
                                           "UMAG_03394"))
dat5 <- melt(dat4, id.vars = "Gene")
dat5$replicate <- sapply(strsplit(as.character(dat5$variable), split = "_"), function(x) x[2])
dat5$time <- sapply(strsplit(as.character(dat5$variable), split = "_"), function(x) x[1])
dat5$time <- ordered(dat5$time, levels = c("axenic", "hpi12", "hpi24", "dpi2", "dpi4", "dpi6", "dpi8", "dpi12"))

dat6 <- dat5
dat6$variable <- NULL
dat6$replicate <- NULL
dat6 <- ddply(.data = dat6, .variables = c("Gene", "time"), .fun = summarize, MeanExpression = mean(value))
dat7 <- dcast(data = dat6, formula = Gene~time)
row.names(dat7) <- dat7$Gene
dat7$Gene <- NULL
labels <- c()
for (i in row.names(dat7)) {
  labels <- c(labels, eval(substitute(expression(italic(x)), list(x = i))))
}

library(gplots)
X11() #Otherwise does not work in RStudio because of non-transparent background.
hm <- heatmap.2(as.matrix(sqrt(dat7)),
        distfun = function(x) dist(x, method = "canberra"),
        hclustfun = function(d) hclust(d, method = "average"),
        reorderfun = function(x, w) return(rev(reorder(x, w))),
        dendrogram = "row",
        Colv = NA, scale = "none",
        labRow = labels,
        col = brewer.pal(9, "Blues"), #rev(heat.colors(50)),
        margins = c(5,15),
        lmat=rbind( c(0, 3), c(2,1), c(4,4)),
        lhei=c(0.5, 4, 1),
        trace = "none",
        density.info = "none",
        key.title = NA,
        key.xlab = "Expression level (normalized read count)",
        key.ylab = NA,
        key.par = list(mar = c(5,1,1,2)),
        key.xtickfun=function() {
          breaks <- parent.frame()$breaks
          return(list(
            at=parent.frame()$scale01(breaks),
            labels=c(as.character(round(breaks*breaks)))
          ))
        })
p3 <- recordPlot()
dev.off()

# Plot profile, in the same order as the dendrogram:
hm <- heatmap(as.matrix(sqrt(dat7)),
                distfun = function(x) dist(x, method = "canberra"),
                hclustfun = function(d) hclust(d, method = "average"),
                reorderfun = function(x, w) return(rev(reorder(x, w))),
                Colv = NA, scale = "none",
                labRow = labels,
                col = brewer.pal(9, "Blues"), #rev(heat.colors(50)),
                margins = c(5,15),
                keep.dendro = TRUE) #Note, heatmap.2 does not keep the dendrogram

dat2 <- subset(dat, Gene %in% c("UMAG_04486", "UMAG_06506", "UMAG_03869", "UMAG_10980", "UMAG_06474", "UMAG_10585", "UMAG_04094", "UMAG_04308", "UMAG_06476", "UMAG_12076", "UMAG_05977"))
dat2$Gene <- ordered(dat2$Gene, levels = rev(labels(hm$Rowv)))
dat3 <- melt(dat2, id.vars = "Gene")
dat3$replicate <- sapply(strsplit(as.character(dat3$variable), split = "_"), function(x) x[2])
dat3$time <- sapply(strsplit(as.character(dat3$variable), split = "_"), function(x) x[1])
dat3$time <- ordered(dat3$time, levels = c("axenic", "hpi12", "hpi24", "dpi2", "dpi4", "dpi6", "dpi8", "dpi12"))

# Second look at all recQ
p2 <- ggplot(dat3, aes(x=as.numeric(time), y=value, group = paste(Gene, replicate))) +
  geom_line(alpha = 0.5) + geom_smooth(se = FALSE, aes(group = Gene)) +
  scale_x_continuous(breaks = as.numeric(unique(dat3$time)), labels = levels(dat3$time)) +
  scale_y_sqrt() +
  facet_wrap(~Gene, nrow = 3) +
  ylab("Expression level\n(normalized read count)") +
  xlab("Time") +
  theme_bw() +
  theme(strip.text = element_text(face = "italic"),
        axis.text.x = element_text(angle = 90))
p2


library(cowplot)
library(gridGraphics)
p23 <- plot_grid(p2, p3, nrow = 1, rel_widths = c(2.2,1), labels = c("B", "C"))
p <- plot_grid(p1, p23, nrow = 2, rel_heights = c(1,2), labels =c("A", ""))

save_plot(p, filename = "UMAG_11065-GeneExpression.pdf", base_width = 12, base_height = 9)

# Statistics:

# Are genes more expressed during infection?
m <- aov(value~Gene*as.ordered(time), dat5)
hist(resid(m))
plot(m)
shapiro.test(resid(m))
library(lmtest)
hmctest(m)
dwtest(m)

library(MASS)
bc <- boxcox(m, lambda = seq(0, 1, length.out = 1000))
l <- bc$x[which.max(bc$y)]
m2 <- aov((value^l - 1)/l~Gene*time, dat5)
hist(resid(m2))
plot(m2)
shapiro.test(resid(m2))
hmctest(m2)
dwtest(m2)
anova(m2)

plot(TukeyHSD(m2, which = "time"), las = 2)

# Does the similarity of expression profiles follow the phylogeny?
library(ape)
phy <- read.tree("../um11065_relativesNewGenomeVersion/Guidance/4thPass/MSA-PhyML_tree.rooted.dnd")
phy <- drop.tip(phy, "FOXG_04692")
plot(phy)

epn <- as.phylo(hclust(dist(as.matrix(sqrt(dat7)), method = "canberra")), method = "average")
plot(epn)

comparePhylo(phy, epn)

mantel.test(cophenetic(phy), cophenetic(epn), nperm = 10000)
```
