## Supplementary File S1 for "The insertion of a mitochondrial selfish element into the nuclear genome and its consequences": InfectionResultsAnalysis.R

```
# Created on 28/12/19 by jdutheil

dat <- read.csv("Infection12dpi.csv", header = TRUE)

# Remove columns with no observation:
selection <- names(which(colSums(dat[,-1]) > 0))
dat2 <- dat[,c("Strain", selection)]

# Test homogeneity of replicates:
chisq.test(as.matrix(dat2[1:3, -1]), simulate.p.value = TRUE, B = 1000000) #p-value = 0.347
chisq.test(as.matrix(dat2[4:6, -c(1,7)]), simulate.p.value = TRUE, B = 1000000) #p-value = 0.829

# We average per strain:
dat3 <- rbind(colSums(dat2[1:3, -1]), colSums(dat2[4:6, -1]))
chisq.test(dat3, simulate.p.value = TRUE, B = 1000000) # p-value = 0.454


# Plot:
dat4 <- read.csv("Infection12dpi.csv", header = TRUE)
dat4 <- dat4[,-6] # Remove "single tumor", are this was never observed, but keep "no symptoms"
dat4$Replicate <- c(1,2,3,1,2,3)

library(plyr)
dat5 <- ddply(.data = dat4, .variables = c("Strain", "Replicate"), .fun = function(x) {
  x <- x[, ! names(x) %in% c("Strain", "Replicate")]
  return(x / sum(x))
})

counts <- data.frame(Strain = c(rep("SG200", 3), rep("SG200~Delta~11064~Delta~11065", 3)),
                     Replicate = c(1,2,3,1,2,3),
                     Count = paste("n =", rowSums(dat4[,-c(1,9)])))

library(reshape2)
dat6 <- dat5
names(dat6) <- gsub(names(dat6), pattern = "\\.", replacement = " ")
dat6$Strain <- c(rep("SG200", 3), rep("SG200~Delta~11064~Delta~11065", 3))
dat6 <- melt(dat6, id.vars = c("Strain", "Replicate"), variable.name = "Symptom", value.name = "Proportion")

library(ggplot2)
library(RColorBrewer)
colors <- brewer.pal(6, "Dark2")
p <- ggplot() + 
  geom_col(data = dat6, aes(x = Replicate, y = Proportion, fill = Symptom), col = "black") +
  facet_grid(~Strain, labeller = label_parsed) +
  scale_fill_manual(values = c("no symptomes" = "white",
                               "chlorosis" = colors[1],
                               "ligula swelling" = colors[2],
                               "small tumors" = colors[3],
                               "tumors" = colors[4],
                               "heavy" = colors[5],
                               "dead plants" = colors[6])) +
  geom_text(data = counts, aes(x = Replicate, y = 1.05, label = Count), size = 3) +
  theme_bw() 
p

# A more conventional picture:

dat4 <- read.csv("Infection12dpi.csv", header = TRUE)
dat4 <- dat4[,-6] # Remove "single tumor", are this was never observed, but keep "no symptoms"

library(plyr)
dat4 <- ddply(.data = dat4, .variables = "Strain", .fun = function(x) {
  x <- x[, names(x) != "Strain"]
  return(colSums(x))
})

dat5 <- ddply(.data = dat4, .variables = "Strain", .fun = function(x) {
  x <- x[, names(x) != "Strain"]
  return(x / sum(x))
})

counts <- data.frame(Strain = c("SG200", "Deletion"),
                     Count = paste("n =", rowSums(dat4[,-c(1,9)])))

library(reshape2)
dat6 <- dat5
names(dat6) <- gsub(names(dat6), pattern = "\\.", replacement = " ")
dat6$Strain <- factor(c("SG200", "Deletion"), levels = c("SG200", "Deletion"))
dat6 <- melt(dat6, id.vars = "Strain", variable.name = "Symptom", value.name = "Proportion")

library(ggplot2)
library(RColorBrewer)
library(scales)
colors <- brewer.pal(6, "YlOrBr")
p <- ggplot() + 
  geom_col(data = dat6, aes(x = Strain, y = Proportion, fill = Symptom), col = "black", width = 0.6) +
  scale_x_discrete(labels = c("SG200" = "SG200", "Deletion" = parse(text = "SG200~Delta~11064~Delta~11065"))) +
  scale_y_continuous(labels = scales::percent) +
  scale_fill_manual(values = c("no symptomes" = "white",
                               "chlorosis" = colors[1],
                               "ligula swelling" = colors[2],
                               "small tumors" = colors[3],
                               "tumors" = colors[4],
                               "heavy" = colors[5],
                               "dead plants" = colors[6])) +
  geom_text(data = counts, aes(x = Strain, y = 1.05, label = Count), size = 3) +
  ylab("Symptoms of infected plants in %")
p


ggsave(p, filename = "Infection12dpi.svg", width = 5, height = 5)
```
