## Supplementary File S1 for "The insertion of a mitochondrial selfish element into the nuclear genome and its consequences": LICENSE

                   GNU LESSER GENERAL PUBLIC LICENSE
                       Version 3, 29 June 2007

 Copyright (C) 2007 Free Software Foundation, Inc. <http://fsf.org/>
 Everyone is permitted to copy and distribute verbatim copies
 of this license document, but changing it is not allowed.


  This version of the GNU Lesser General Public License incorporates
the terms and conditions of version 3 of the GNU General Public
License, supplemented by the additional permissions listed below.

  0. Additional Definitions.

  As used herein, "this License" refers to version 3 of the GNU Lesser
General Public License, and the "GNU GPL" refers to version 3 of the GNU
General Public License.

  "The Library" refers to a covered work governed by this License,
other than an Application or a Combined Work as defined below.

  An "Application" is any work that makes use of an interface provided
by the Library, but which is not otherwise based on the Library.
Defining a subclass of a class defined by the Library is deemed a mode
of using an interface provided by the Library.

  A "Combined Work" is a work produced by combining or linking an
Application with the Library.  The particular version of the Library
with which the Combined Work was made is also called the "Linked
Version".

  The "Minimal Corresponding Source" for a Combined Work means the
Corresponding Source for the Combined Work, excluding any source code
for portions of the Combined Work that, considered in isolation, are
based on the Application, and not on the Linked Version.

  The "Corresponding Application Code" for a Combined Work means the
object code and/or source code for the Application, including any data
and utility programs needed for reproducing the Combined Work from the
Application, but excluding the System Libraries of the Combined Work.

  1. Exception to Section 3 of the GNU GPL.

  You may convey a covered work under sections 3 and 4 of this License
without being bound by section 3 of the GNU GPL.

  2. Conveying Modified Versions.

  If you modify a copy of the Library, and, in your modifications, a
facility refers to a function or data to be supplied by an Application
that uses the facility (other than as an argument passed when the
facility is invoked), then you may convey a copy of the modified
version:

   a) under this License, provided that you make a good faith effort to
   ensure that, in the event an Application does not supply the
   function or data, the facility still operates, and performs
   whatever part of its purpose remains meaningful, or

   b) under the GNU GPL, with none of the additional permissions of
   this License applicable to that copy.

  3. Object Code Incorporating Material from Library Header Files.

  The object code form of an Application may incorporate material from
a header file that is part of the Library.  You may convey such object
code under terms of your choice, provided that, if the incorporated
material is not limited to numerical parameters, data structure
layouts and accessors, or small macros, inline functions and templates
(ten or fewer lines in length), you do both of the following:

   a) Give prominent notice with each copy of the object code that the
   Library is used in it and that the Library and its use are
   covered by this License.

   b) Accompany the object code with a copy of the GNU GPL and this license
   document.

  4. Combined Works.

  You may convey a Combined Work under terms of your choice that,
taken together, effectively do not restrict modification of the
portions of the Library contained in the Combined Work and reverse
engineering for debugging such modifications, if you also do each of
the following:

   a) Give prominent notice with each copy of the Combined Work that
   the Library is used in it and that the Library and its use are
   covered by this License.

   b) Accompany the Combined Work with a copy of the GNU GPL and this license
   document.

   c) For a Combined Work that displays copyright notices during
   execution, include the copyright notice for the Library among
   these notices, as well as a reference directing the user to the
   copies of the GNU GPL and this license document.

   d) Do one of the following:

       0) Convey the Minimal Corresponding Source under the terms of this
       License, and the Corresponding Application Code in a form
       suitable for, and under terms that permit, the user to
       recombine or relink the Application with a modified version of
       the Linked Version to produce a modified Combined Work, in the
       manner specified by section 6 of the GNU GPL for conveying
       Corresponding Source.

       1) Use a suitable shared library mechanism for linking with the
       Library.  A suitable mechanism is one that (a) uses at run time
       a copy of the Library already present on the user's computer
       system, and (b) will operate properly with a modified version
       of the Library that is interface-compatible with the Linked
       Version.

   e) Provide Installation Information, but only if you would otherwise
   be required to provide such information under section 6 of the
   GNU GPL, and only to the extent that such information is
   necessary to install and execute a modified version of the
   Combined Work produced by recombining or relinking the
   Application with a modified version of the Linked Version. (If
   you use option 4d0, the Installation Information must accompany
   the Minimal Corresponding Source and Corresponding Application
   Code. If you use option 4d1, you must provide the Installation
   Information in the manner specified by section 6 of the GNU GPL
   for conveying Corresponding Source.)

  5. Combined Libraries.

  You may place library facilities that are a work based on the
Library side by side in a single library together with other library
facilities that are not Applications and are not covered by this
License, and convey such a combined library under terms of your
choice, if you do both of the following:

   a) Accompany the combined library with a copy of the same work based
   on the Library, uncombined with any other library facilities,
   conveyed under the terms of this License.

   b) Give prominent notice with the combined library that part of it
   is a work based on the Library, and explaining where to find the
   accompanying uncombined form of the same work.

  6. Revised Versions of the GNU Lesser General Public License.

  The Free Software Foundation may publish revised and/or new versions
of the GNU Lesser General Public License from time to time. Such new
versions will be similar in spirit to the present version, but may
differ in detail to address new problems or concerns.

  Each version is given a distinguishing version number. If the
Library as you received it specifies that a certain numbered version
of the GNU Lesser General Public License "or any later version"
applies to it, you have the option of following the terms and
conditions either of that published version or of any later version
published by the Free Software Foundation. If the Library as you
received it does not specify a version number of the GNU Lesser
General Public License, you may choose any version of the GNU Lesser
General Public License ever published by the Free Software Foundation.

  If the Library as you received it specifies that a proxy can decide
whether future versions of the GNU Lesser General Public License shall
apply, that proxy's public statement of acceptance of any version is
permanent authorization for you to choose that version for the
Library.
