## Supplementary File S1 for "The insertion of a mitochondrial selfish element into the nuclear genome and its consequences": README.md

Created on 08/01/20 by jdutheil

Retrieve DNA sequences of selected UMAG_11065 relatives:

```R
library(seqinr)
dna <- read.fasta("../all.dna.formated.fa")
pro <- read.fasta("../Guidance/4thPass/MSA.MAFFT.aln.With_Names")
sel <- dna[names(pro)]
write.fasta(sequences=sel, names=names(sel), file.out="UMAG_11065relatives.fasta")
```

Align sequences using Macse:
```r
java -Xmx600m -jar ~/.local/bin/macse_v1.2.jar -prog alignSequences -seq UMAG_11065relatives.fasta

```

PhyML tree (LG+Gamma 4, BEST), after manually selecting sites and sequences that match UMAG_11065.

Convert sequence to PAML format:
```bash
bppseqman param=SeqMan.bpp
```

Run PAML:
```bash
codeml
```

UMAG_11065 seems to be under purifying selection, but the branch below seems to be under positive selection.
We just check with a branchsite model:

```bash
codeml codeml2.ctl
```
The model fails to estimate omega2.
