## Supplementary File S1 for "The insertion of a mitochondrial selfish element into the nuclear genome and its consequences": figTree.R

```
# Created on 16/08/19 by jdutheil
# Plot UMAG_11065 phylogenetic tree

library(ape)

tree <- read.tree("Guidance/4thPass/MSA-PhyML_tree.unrooted.dnd")
tree$node.label <- as.numeric(tree$node.label) * 100
cols <- heat.colors(101, rev = TRUE, )
plot(tree, underscore = TRUE, no.margin = TRUE, align.tip.label = TRUE, edge.width = 2)
nodelabels(tree$node.label, bg = cols[tree$node.label + 1], frame = "c")
add.scale.bar(x = 0, y = 14)

dev.print(pdf, file = "UMAG_11065_tree.pdf", width = 8, height = 6)

# V2: collapsing node and using ggtree:
library(ggtree)

tree <- read.tree("Guidance/4thPass/MSA-PhyML_tree.unrooted.collapsed.dnd")
tree$node.label <- as.numeric(tree$node.label) * 100

p <- ggplot(tree, aes(x, y))
p <- p + geom_tree() + geom_tiplab(align = TRUE) + geom_nodelab(geom = "label", hjust = 0) + geom_treescale()
p <- p + theme_tree() + xlim(0,1) + theme()
p

ggsave(p, filename = "UMAG_11065_tree.pdf", width = 8, height = 6)
```
