## Supplementary File S1 for "The insertion of a mitochondrial selfish element into the nuclear genome and its consequences": formatSeq.R

```
# Created on 27/12/16 by jdutheil
# Rename sequences from PEDANT for easier use with phylogenetic software

library(seqinr)

seq <- read.fasta("all.prot.fa")
l <- names(seq)
l <- sapply(strsplit(l, "@"), function(x) x[2])
write.fasta(seq, names = l, file.out = "all.prot.formated.fa")

seq <- read.fasta("all.dna.fa")
l <- names(seq)
l <- sapply(strsplit(l, "@"), function(x) x[2])
write.fasta(seq, names = l, file.out = "all.dna.formated.fa")
```
