## Supplementary File S1 for "The insertion of a mitochondrial selfish element into the nuclear genome and its consequences": README.md

# Created on 26/12/16 by jdutheil
# Updated data downloaded from PEDANT3

```bash
rm all.dna.fa
rm all.prot.fa
cat *.dna.fa > all.dna.fa
cat *.prot.fa > all.prot.fa
```

To get nicer sequence names:
```bash
Rscript formatSeq.R
```

Plot the tree:
```bash
Rscript figTree.R
```
