## Supplementary File S1 for "The insertion of a mitochondrial selfish element into the nuclear genome and its consequences": README.md

created on 02/02/14 by jdutheil
Uses boxshade to produce xfig file and edit

- Original macse alignment:
RegionChr09+um11064+um11065+abI-AbiIII-P+SrCox1Intron1+TilletiaCox1Intron1_macse_NT-edited.mase

- Edit in seaview, remove UMAG_11065 and UMAG_11064, convert to proteins, remove gap columns, select interesting region and save to Alignment.mase
seaview ~/Data/Fungi/Genomes/MitoTransfer/LookingUpstream/RegionChr09+um11064+um11065+abI-AbiIII-P+SrCox1Intron1+TilletiaCox1Intron1_macse_NT-edited.mase

```bash
boxshade -in=Alignment.aln -def -par=boxshade.par -dev=c -out=Alignment.fig -unix
```

- Open in xfig and export to SVG

- Make further cosmetc changes in inkscape

# Phylogeny

- Make manual selection of sites in seaview (selection is saved in mase file)
- PhyML tree with LG + Gamma(4), BEST. save rooted tree.
